# Pyruvate dehydrogenase kinase supports macrophage NLRP3 inflammasome activation during acute inflammation

**DOI:** 10.1101/2021.10.02.462869

**Authors:** Allison K. Meyers, Zhan Wang, Wenzheng Han, Qingxia Zhao, Manal Zabalawi, Likun Duan, Juan Liu, Qianyi Zhang, Rajesh K Manne, Felipe Lorenzo, Matthew A. Quinn, Qianqian Song, Daping Fan, Hui-Kuan Lin, Cristina M. Furdui, Jason W. Locasale, Charles E. McCall, Xuewei Zhu

## Abstract

Activating macrophage NLRP3 inflammasome can promote excessive inflammation, with severe cell and tissue damage and organ dysfunction. Here, we show that pharmacological or genetic inhibition of pyruvate dehydrogenase kinase (PDHK) significantly attenuates NLRP3 inflammasome activation in murine and human macrophages and septic mice by lowering caspase-1 cleavage and IL-1β secretion. Inhibiting PDHK reverses NLRP3 inflammasome-induced metabolic reprogramming, enhances autophagy, promotes mitochondrial fusion over fission, preserves cristae ultrastructure, and attenuates mitochondrial ROS production. The suppressive effect of PDHK inhibition on the NLRP3 inflammasome is independent of its canonical role as a pyruvate dehydrogenase regulator. We suggest that PDHK inhibition improves mitochondrial fitness by reversing NLRP3 inflammasome activation in acutely inflamed macrophages.

## Introduction

In sensing danger and promoting inflammation from diverse environmental threats, macrophages express pattern recognition toll-like receptors (TLRs) and intracellular NOD-like receptors, such as NOD-, LRP-, and pyrin domain-containing protein 3 (NLRP3). Activation of NLRP3 results in the assembly of the NLRP3 inflammasome, leading to self-cleavage of caspase-1, processing of IL-1β and IL-18, and pyroptotic cell death (pyroptosis).^1-6^ Macrophage NLRP3 inflammasomes play a vital role in host defense against invading microbes.^7,8^ Exaggerated NLRP3 inflammasome activation increases tissue inflammation, cell death, and organ damage. Moreover, pyroptosis can reduce functional monocyte/macrophage numbers and activate coagulation by releasing tissue factors.^9^ Uncontrolled NLRP3 inflammasome activation accelerates the pathogenesis of many inflammatory diseases, including cardiovascular dysfunction, type II diabetes, sepsis, and COVID-19 syndrome.^10-14^ Uncovering key regulators of NLRP3 inflammasome activation may lead to therapeutic strategies for preventing and treating inflammatory diseases.

Mitochondria inform NLRP3 inflammasome activation. Dysfunctional mitochondria release mitochondrial reactive oxygen species (mtROS),^15,16^ newly synthesized mitochondrial DNA,^17,18^ and expose cardiolipin on the outer membrane to activate the NLRP3 inflammasome^19^. Mitochondrial TCA cycle intermediates succinate,^20,21^ itaconate,^22-24^ and fumarate^25^ differentially regulate NLRP3 inflammasome activation. Very recently, mitochondrial ATP instead of ROS has been proposed to activate the NLRP3 inflammasome,^26,27^ complicating the role of mitochondria in regulating the NLRP3 inflammasome.

Mitochondrial pyruvate dehydrogenase (PDH) complex (PDC) is the gate-keeping and rate-limiting enzyme for mitochondrial glucose oxidation by decarboxylating pyruvate to increase acetyl-CoA.^28^ Pyruvate dehydrogenase kinase (PDHK) isoforms 1-4 reversibly phosphorylate the PDHe1α subunit to inhibit PDC, whereas pyruvate dehydrogenase phosphatase (PDP) dephosphorylates and activates PDC.^28-31^ Pyruvate dehydrogenase kinase 1 (PDHK1) is persistently expressed in monocytes and macrophages in septic mice and human sepsis blood monocytes.^32^ Targeting PDHK with its prototypic inhibitor dichloroacetate (DCA), a structural analog of pyruvate (the endogenous inhibitor of PDHK),^28^ protects mice against cecal-ligation and puncture (CLP)-induced polymicrobial infection.^33^ However, the molecular mechanisms underlying DCA-induced septic protection are unclear.

The present study shows that pharmacological or genetic inhibition of PDHK suppresses macrophage NLRP3 inflammasome activation and promotes macrophage metabolic and mitochondrial homeostasis during NLRP3 inflammasome activation, using *in vitro* sepsis cell modeling and septic mice. Unexpectedly, the suppressive effect of PDHK inhibition on the NLRP3 inflammasome is independent of pyruvate dehydrogenase, autophagy, or ROS production. Thus, this study uncovers a non-canonical role for mitochondrial PDHK in promoting mitochondrial stress and activating NLRP3 inflammasome during acute inflammation.

## Results

### PDHK inhibition lowers macrophage NLRP3 inflammasome activation

To examine the effect of PDHK inhibition on macrophage NLRP3 inflammasome activation, we utilized two PDHK pharmacological inhibitors: DCA and JX06. DCA blocks all four PDHK isoforms by binding to the pyruvate kinase pocket. In contrast, JX06 preferentially blocks PDHK isoform 1 by forming a disulfide bond with Cys240 within the ATP pocket.^34^ We treated murine bone marrow-derived macrophages (BMDMs) with DCA or JX06 for 30 min before LPS priming or after LPS priming, by adding the inhibitors with the NLRP3 inflammasome inducers ATP, nigericin, and monosodium urate crystals (MSU) (**Figure 1A**). JX06 significantly reduced ATP, nigericin, or MSU-induced IL-1β secretion regardless of being added before or after LPS priming, but DCA inhibited MSU-induced IL-1β secretion only when added after the LPS priming step (**Figures 1B-1C**). Both DCA and JX06 more profoundly repressed IL-1β secretion when added after LPS priming (**Figures 1B-1C**). Therefore, in subsequent experiments, DCA or JX06 treatment followed LPS priming unless specifically indicated. Both DCA and JX06 dose-dependently suppressed ATP-induced IL-1β secretion (**Figures 1D-1E**), IL-1β cleavage (IL-1β p17), and caspase-1 cleavage (caspase-1 p20) (**Figures 1F-1G**) without altering intracellular protein levels of NLRP3 inflammasome components (**Figures 1F-1G**). Consistently, DCA or JX06 showed a minor to no effect on the transcript expression of pro-IL-1β or NLRP3 when added before or after LPS priming (**Figures 1H-1I**). Additionally, DCA and JX06 reduced caspase-1 cleavage in macrophages treated with LPS plus nigericin (**Figures S1A-S1B**) or MSU (**Figures S1C-S1D**). These results suggest that PDHK inhibition blocks macrophage NLRP3 inflammasome activation primarily by attenuating caspase-1 cleavage (Signal 2) but not LPS priming (Signal 1). Caspase-1 cleaves gasdermin D (GSDMD), leading to plasma membrane rupture, cell swelling (ballooning effect), and pyroptotic death.^35-37^ PDHK inhibition prevented NLRP3-induced cellular swelling in LPS plus ATP-treated macrophages (**Figures S1E-S1F**). PDHK inhibition also protected macrophages against LPS plus ATP-induced pyroptotic death, as shown by decreased propidium iodide (PI) or 7-amino-actinomycin D (7-AAD) staining (**Figures 1J-1K**) and less full-length and cleaved N-terminal GSDMD in the culture supernatant (**Figure 1L**).

**Figure 1.**
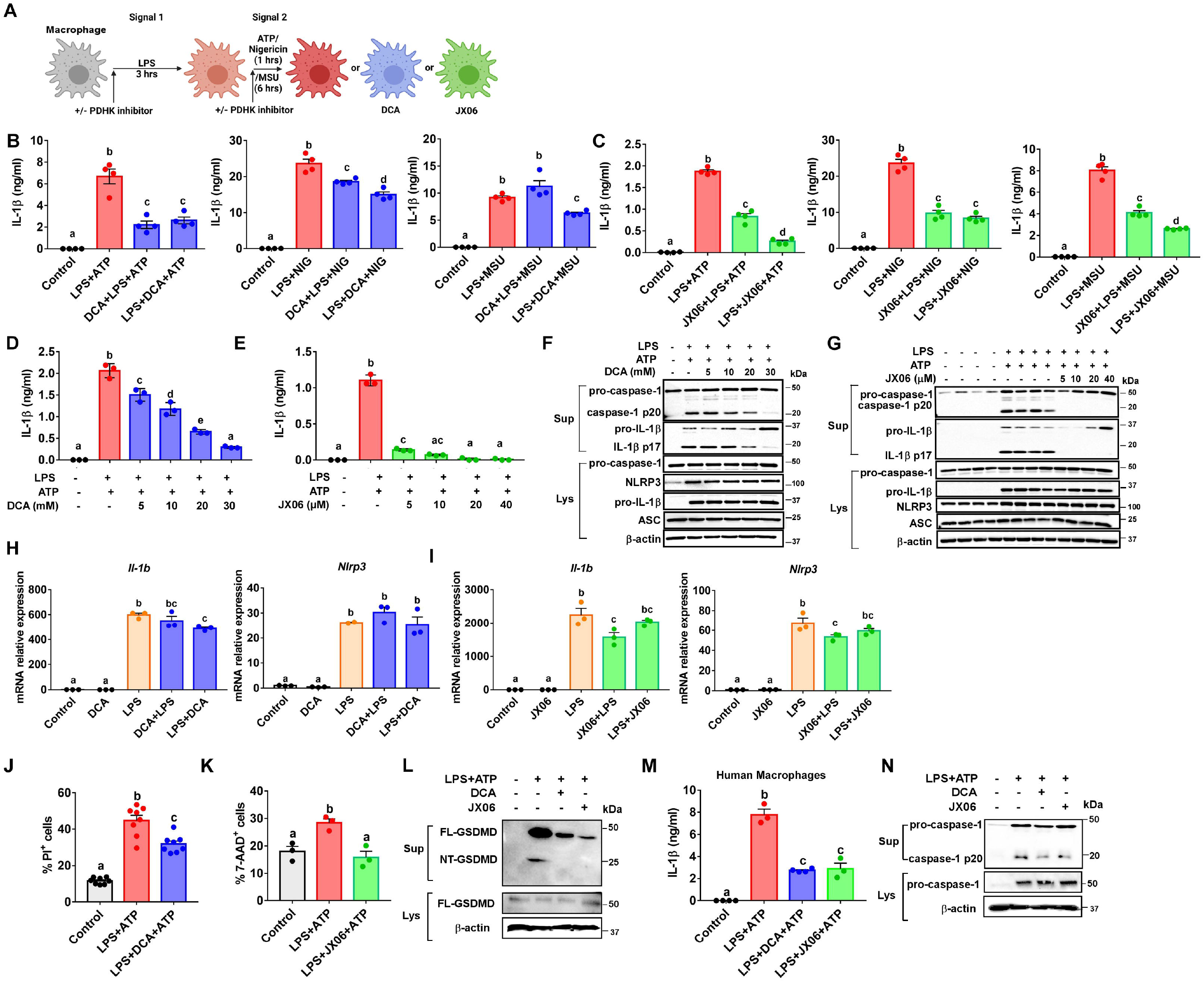
PDHK inhibition lowers macrophage NLRP3 inflammasome activation and cell death. (**A**) Schematic of macrophage treatments and timing. (**B-C**) IL-1β concentrations in the culture supernatant were analyzed by ELISA. 20 mM DCA, 10 µM JX06, or DMSO (vehicle control for JX06) were added before or after LPS priming. (**D-E**) IL-1β concentrations in the culture supernatant were analyzed by ELISA. DCA, JX06, or DMSO (vehicle control for JX06) were added after LPS priming. (**F-G**) Protein expression in the culture supernatant (Sup) and whole-cell lysates (Lys) was analyzed by immunoblotting. DCA, JX06, or DMSO (vehicle control for JX06) were added after LPS priming. (**H-I**) mRNA expression of NLRP3 inflammasome components was measured by qPCR. 20 mM DCA or 10 µM JX06 were added before or after LPS priming. (**J-K**) Cell death was measured by staining macrophages with 7-AAD or PI and analyzed by flow cytometry. (**L**) Full length (FL) and N-terminal (NT) GSDMD protein expression in the culture supernatant (Sup) and whole-cell lysates (Lys) was analyzed by immunoblotting. (**M-N**) IL-1β concentrations in the culture supernatant were analyzed by ELISA and protein expression in the culture supernatant (Sup) and whole-cell lysates (Lys) was analyzed by immunoblotting. 20 mM DCA or 10 µM JX06 were added after LPS priming and together with ATP in human macrophages. Data are reprehensive of three independent experiments (n=3-4 samples per group). Groups with different letters are significantly different (p<0.05), one-way ANOVA with post hoc Tukey’s multiple comparisons test.

DCA and JX06 also lowered LPS plus ATP-induced IL-1β secretion and caspase-1 cleavage in human peripheral blood mononuclear cells (PBMCs)-derived macrophages (**Figures 1M-1N**). In contrast, DCA or JX06 showed no effect on AIM2 inflammasome-mediated IL-1β production (**Figure S1G**) and did not reduce TNF or IL-6 secretion regardless of being added before or after LPS treatment (**Figures S1H-S1I**). Unlike DCA, JX06 treatment moderately lowered LPS-induced TNF or IL-6 production, when added before LPS stimulation (**Figures S1H-S1I**). These results suggest that PDHK inhibition has a more profound inhibitory effect on NLRP3 inflammasome-mediated inflammation.

### PDHK inhibition rewires cellular metabolism in NLRP3 inflammasome-activated macrophages

We next analyzed unbiased metabolomics with or without PDHK inhibition following NLRP3 inflammasome activation in macrophages. A principal component analysis (PCA) of cellular metabolites (**Figure 2A**) demonstrated the distinct segregation of each treatment group. Among the 236 metabolites detected, 101 metabolites (red) showed significant accumulation in the inflammasome-activated vs. control macrophages (**Figure 2B**). The predominant downregulated metabolites (green, n=93) in the inflammasome-activated vs. control macrophages were antioxidants (**Figure 2B**). Metabolite enrichment analysis revealed broad metabolic reprogramming in macrophages during NLRP3 inflammasome activation (**Figure 2C**) with JX06 significantly altering 119 metabolites (**Figure 2D**). JX06 treatment prominently up-regulated bilirubin in LPS plus ATP-treated cells. Other metabolites identified in metabolic pathway analysis included porphyrin and chlorophyll, arachidonic acid, pyruvate, and TCA cycle intermediates (**Figures 2E-2F**). Collectively, these results indicate that PDHK inhibition by JX06 broadly influences cellular lipid, glucose, redox, and TCA cycle metabolism, demonstrating that PDHK is a vital kinase that controls macrophage metabolism under conditions of the NLRP3 inflammasome activation.

**Figure 2.**
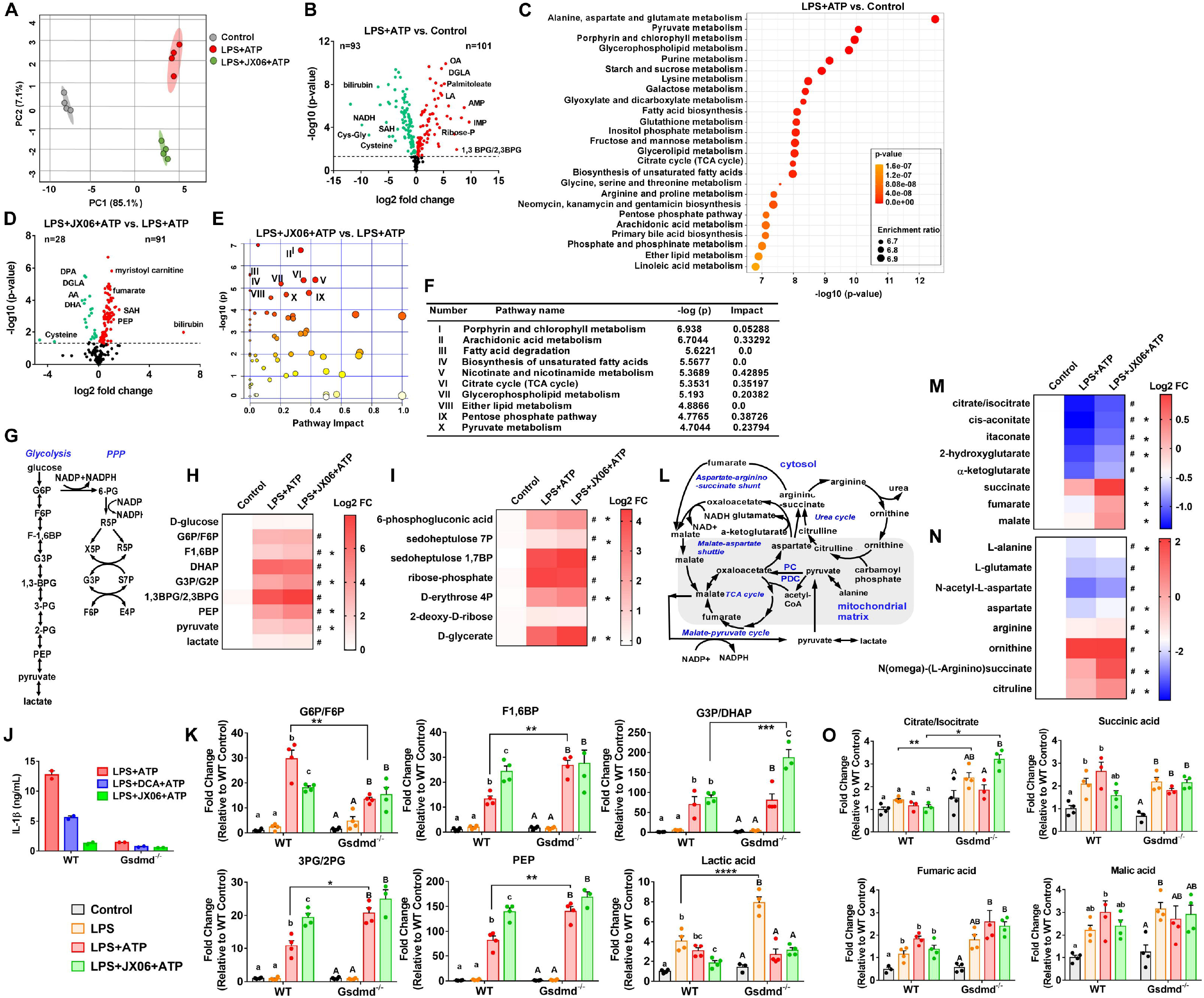
PDHK inhibition rewires cellular metabolism in NLRP3 inflammasome-activated macrophages. (**A**) Principal component analysis of the BMDM metabolites from untargeted metabolomics data (4 technical replicates per group). (**B**) Volcano plot of the metabolites in LPS+ATP vs. control macrophages, plotted using GraphPad 7.0. (**C**) The top 25 enriched metabolic pathways in LPS+ATP treated vs. control BMDMs, identified by enrichment analysis and ranked according to p values. (**D**) Volcano plot of the metabolites in JX06 vs. LPS+ATP treated macrophages, plotted using GraphPad 7.0. (**E-F**) (**E-F**) The top 10 metabolic pathways altered by JX06 treatment, identified by the pathway analysis and ranked according to p values. (**G**) Diagram of glycolysis and PPP. (**H**) Heatmap of the intermediates in the glycolysis pathway from the untargeted metabolomics analysis. (**I**) Heatmap of the intermediates in PPP from the untargeted metabolomics analysis. (**J**) IL-1β concentrations in the culture supernatant from WT and Gsdmd^-/-^ mouse BMDMs treated with LPS plus ATP in the presence or absence of DCA or JX06. (**K**) Cellular glycolytic intermediates in WT and Gsdmd^-/-^ mouse BMDMs (J) were quantified by targeted metabolomics (4 technical replicates per group). (**L**) Diagram of the TCA cycle and its coupling pathways. (**M**) Heatmap of the intermediates in the TCA cycle from the untargeted metabolomics analysis. (**N**) Heatmap of the intermediates in the PC-mediated anaplerosis pathway, aspartate-argininosuccinate shunt, and urea cycle from the untargeted metabolomics analysis. (**O**) Cellular TCA cycle intermediates in WT and Gsdmd^-/-^ mouse BMDMs (J) were quantified by targeted metabolomics. ^#^: p<0.05, between LPS+ATP and control groups (Figures 2H, 2I, 2M, and 2N); *: p<0.05, between LPS+ATP and LPS+JX06+ATP groups (Figures 2H, 2I, 2M, and 2N); one-way ANOVA with post hoc Tukey’s multiple comparisons test. * p<0.05, ** p<0.01. Different lower case letters are used to denote significant differences (p<0.05) among treatments or treatments in WT cells (Figures 2K and 2O). Uppercase letters are used to denote significant differences (p<0.05) among groups in Gsdmd^-/-^ cells (Figures 2K and 2O). Two-way ANOVA with post hoc Tukey’s multiple comparisons test.

Pyruvate metabolism and PPP (**Figure 2G**) are among top metabolic pathways altered by JX06 treatment (**Figure 2F**). LPS plus ATP significantly increased glycolytic intermediate levels in macrophages, as measured by our untargeted metabolomics analysis (**Figure 2H)**. JX06 treatment further increased glycolytic intermediate accumulation except for intracellular lactate (**Figure 2H**). JX06 also increased the levels of PPP intermediates (**Figure 2I**). Given that pyroptosis induces plasma membrane rupture and that PDHK inhibition dampens pyroptotic cell death (**Figures 1J-1L**), we next examined the impact of pyroptosis on glycolytic metabolites using GSDMD-deficient macrophages via targeted metabolomics analysis. GSDMD-deficient macrophages markedly reduced IL-1β secretion in response to LPS plus ATP stimulation (**Figure 2J**). We did not observe a significant increase in glycolytic metabolites except for lactic acid in both LPS-only treated wild-type (WT) and GSDMD-deficient macrophages (**Figure 2K**). Consistent with untargeted metabolomics analysis, LPS plus ATP increased glycolytic metabolites compared to the control and JX06 accentuated these metabolites in WT cells except for downregulated G6P/F6P (**Figure 2K**). Also observed were down-regulated G6P/F6P but up-regulated F1,6BP, 3PG/2PG, and PEP in LPS plus ATP-treated GSDMD-deficient vs. WT macrophages (**Figure 2K**). These results suggest that pyroptosis alters glycolytic metabolite concentrations in NLRP3 inflammasome-activated macrophages but does not increase glycolytic metabolites universally. In response to JX06 treatment, GSDMD-deficient macrophages showed a significant increase in G3P/DHAP but only a trend toward an increase in 3PG/2PG and PEP relative to WT (**Figure 2K**). Together, the data suggest that macrophages reprogram glucose metabolism in response to the NLRP3 inflammasome activation and during PDHK inhibition.

Next assessed was whether PHK inhibition alters the TCA cycle (**Figure 2L**). NLRP3 inflammasome activation by LPS+ATP significantly lowered TCA cycle intermediates citrate/isocitrate and cis-aconitate, but increased succinate relative to untreated control cells (**Figure 2M**). Concurrent with the changes to TCA cycle intermediates, NLRP3 inflammasome activation significantly lowered the levels of aspartate and alanine, a pyruvate carboxylase-dependent anaplerotic substrate and increased urea cycle intermediates such as ornithine and citrulline (**Figure 2N**). Inhibition of PDHK by JX06 in LPS plus ATP-treated macrophages increased TCA cycle intermediates (**Figure 2M**). Also increased were aspartate, alanine, argininosuccinate, arginine, and citrulline (**Figure 2N**). Distinct from its changes in glycolytic metabolites, GSDMD deficiency showed a minor effect on the TCA cycle metabolites in LPS- or LPS plus ATP-treated macrophages (**Figure 2O**), with or without PDHK inhibition. JX06 also showed a minimal impact on the TCA cycle metabolites in LPS plus ATP-treated WT and Gsdmd^-/-^ macrophages, suggesting that the metabolic changes in PDHK-inhibited macrophages are not solely attributable to the difference in plasma membrane integrity.

### PDHK inhibition rescues metabolic paralysis in NLRP3 inflammasome-activated macrophages

Given that the metabolomics analysis only provides a snapshot of the concentrations of cellular metabolites, we next performed a ^13^C_6_ glucose stable isotope tracing experiment to assess the effect of PDHK inhibition on glucose utilization in macrophages during acute inflammation (**Figure 3A**). Macrophages were labeled with [U-^13^C]-glucose for 90 min post ATP±JX06 treatment (**Figure 3B**), achieving 95% isotopic enrichment of cellular M+6 labeled glucose (**Figure 3C**). Macrophages enriched ^13^C_6_ glucose-derived carbon in response to inflammatory stimulation and JX06 treatment (**Figure 3D**). Pathway analysis of total ^13^C fraction reveals ^13^C_6_-glucose-enriched metabolites clustered into several pathways in LPS plus ATP vs. LPS only (**Figure 3E**), or vs. JX06 (**Figure 3F**)-treated macrophages. The clusters included glycolysis, pyruvate metabolism, the TCA cycle, and nicotinate and nicotinamide metabolism. LPS-primed macrophages display increased fractional enrichment in most glycolytic intermediates (**Figure 3G**). In contrast, LPS plus ATP-treated macrophages showed overall low fractional enrichment in glycolytic intermediates, suggesting impaired glucose flux through glycolysis. Strikingly, JX06 reversed the defective glycolytic flux (**Figure 3G**) and corrected low glucose flux through PPP in inflammasome-activated cells, as shown by higher abundance in M+5 labeled ribose-5-phosphate and M+7 labeled S7P (**Figure 3H**).

**Figure 3.**
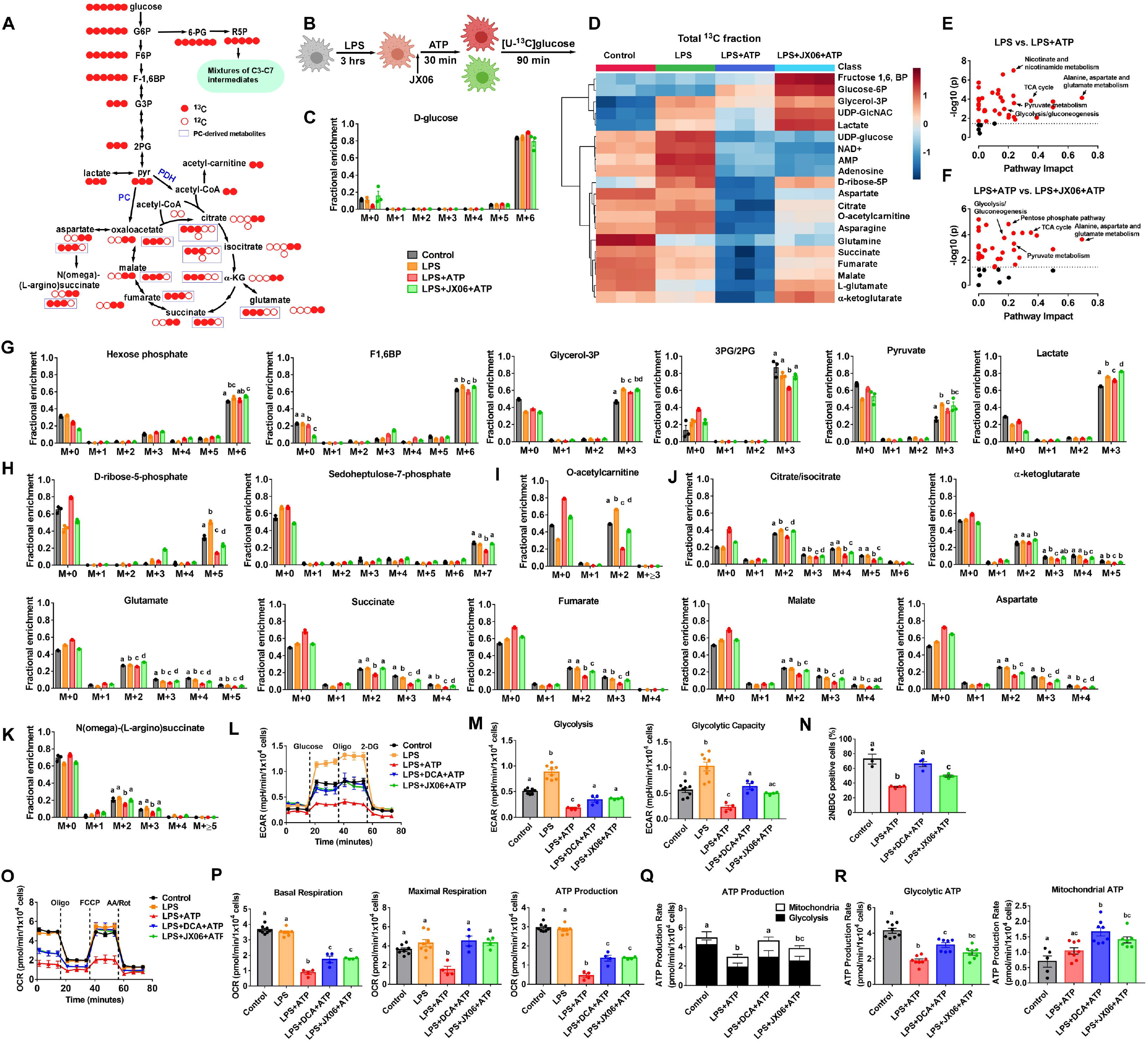
PDHK inhibition improves glucose flux and mitochondrial respiration in NLRP3 inflammasome-activated macrophages. (**A**) Schematic of carbon incorporation into glycolysis, PPP, and the TCA cycle intermediates from [U-^13^C] glucose (red). (**B**) Schematic of *in vitro* [U-^13^C]-glucose labeling protocol. (**C**) Fractional enrichment and labeling patterns of glucose from [U-^13^C] glucose-labeled BMDMs (n=3 technical replicates). Natural abundance correction was performed. (**D**) Heatmap of the top 20 differentially labeled metabolites among groups. Data points reflect the sum of fractional enrichment of all ^13^C isotopologues. (**E-F**) Metabolic pathway analysis of ^13^C-labeled metabolites in BMDMs treated with LPS vs. LPS plus ATP (E) or with LPS+ATP vs. LPS+JX06+ATP (F). (**G-K**) Fractional enrichment and labeling patterns of glycolytic (G), PPP (H), and TCA cycle (I-K) intermediates and/or derivatives from [U-^13^C] glucose-labeled BMDMs. Natural abundance correction was performed. One-way ANOVA analysis was done within a specific ^13^C isotopologue group. (**L-M**) Seahorse analysis of extracellular acidification rates (ECAR) in BMDMs (n= 4-8 technical replicates). (**N**) Glucose uptake in BMDMs treated with LPS+ATP in the presence or absence of DCA, JX06, or 2-NBDG (100 µg/ml) for 30 min. Macrophages were analyzed by flow cytometry (n= 3-4 technical replicates). (**O-P**) Seahorse analysis of oxygen consumption rates (OCR) in BMDMs (n= 4-8 technical replicates). (**Q-R**) Seahorse XF real-time ATP rate analysis of BMDMs. Glycolytic ATP and mitochondrial ATP production rates were calculated (n= 8 technical replicates). Groups with different letters are significantly different (p<0.05); One-way ANOVA with post hoc Tukey’s multiple comparisons test.

^13^C_6_-glucose-derived ^13^C_3_-pyruvate can enter the TCA cycle as citrate with two labeled carbons (M+2) after decarboxylation by PDC or as oxaloacetate with three labeled carbons (M+3) after carboxylation by PC, which then combines with acetyl-CoA to make M+3 citrate (**Figure 3A**) during 1^st^ round TCA cycle. The most abundant isotopologue in the TCA cycle intermediates or their derivatives after 90 min ^13^C_6_-glucose tracing in macrophages was PDH-mediated M+2 carbons (**Figures 3I-3K**). M+3 labeled citrate/isocitrate formed from PC-mediated carboxylation and M+4 and M+≥5 isotopologues in TCA cycle intermediates emerged, perhaps during 2^nd^ round TCA cycle.

LPS alone significantly increased M+2 acetyl-CoA-derived acetylcarnitine (**Figure 3I**) and citrate/isocitrate (**Figure 3J**), suggesting increased ^13^C_3_-pyruvate entering the PDC portal. In contrast, LPS plus ATP-treated macrophages had the lowest fractional enrichment in ^13^C-labeled TCA cycle intermediates, or their derivatives (**Figures 3I-3K**), suggesting inhibited glucose flux into the TCA cycle during NLRP3 inflammasome activation. Notably, JX06 treatment significantly increased the abundance of M+≥2 isotopologues in most TCA cycle metabolites, from both PDC- and PC-mediated pyruvate utilization (**Figures 3I-3K**). These stable isotope labeling data suggest that NLRP3 inflammasome activation reduces glucose flux in macrophages relative to LPS-induced macrophage activation. Significantly, PDHK inhibition rescues the TCA cycle by upregulating PDC- and PC-mediated glucose oxidation.

Consistent with the glucose flux patterns, LPS priming enhanced glycolysis and glycolysis capacity in macrophages, as indicated by a significant increase in extracellular acidification rate (ECAR) after injection of glucose and oligomycin in the respirometer (**Figures 3L-3M**). In contrast, LPS plus ATP significantly lowered glycolysis and glycolysis capacity (**Figures 3L-3M**). DCA and JX06 increased ECAR and restored glycolysis to the control level (**Figures 3L-3M**). However, DCA but not JX06 restored the glycolytic capacity of the LPS+ATP-treated macrophages to the control level (**Figure 3M**). DCA did not impact glycolysis or glycolytic capacity in LPS-only treated macrophages (**Figures S2A-S2B**). LPS plus ATP treatment also significantly impaired macrophage glucose uptake compared to control, while PDHK inhibition increased glucose uptake (**Figure 3N**). Thus, PDHK inhibition promotes glucose uptake and glycolysis in NLRP3 inflammasome-activated macrophages. A Seahorse Mito Stress test showed that LPS plus ATP treatment, but not LPS priming, significantly lowered mitochondrial respiration (**Figures 3O-3P**). PDHK inhibition improved mitochondrial respiration in LPS plus ATP-treated macrophages (**Figure 3P**) but had no impact on mitochondrial respiration in LPS-only treated cells (**Figures S3C-S3D**). Moreover, pretreating macrophages with glycine to block plasma membrane rupture did not affect mitochondrial respiration (**Figures S3E-S3F**). LPS plus ATP treatment also decreased total ATP production in macrophages, as quantified by the Seahorse XF real-time ATP rate assay (**Figure 3Q**). PDHK inhibition by DCA rejuvenated glycolytic and mitochondrial ATP production (**Figure 3R**). A similar trend occurred in JX06-treated macrophages (**Figure 3R**). Thus, PDHK inhibition of NLRP3 inflammasome in macrophages restores macrophage metabolic fueling and energy homeostasis.

### PDHK inhibition-mediated NLRP3 inflammasome inactivation is independent of glucose metabolism or PDH

To assess whether the enhanced glucose utilization accounts for PDHK inhibition-mediated NLRP3 inflammasome inactivation, we blocked glycolysis, PPP, and mitochondrial glucose metabolism by adding pharmacological inhibitors 30 min before ATP ± DCA stimulation (**Figure 4A**). Blockade of glycolysis using 2-DG (blocking hexose kinase), heptelidic acid (HA, blocking GAPDH), or sodium oxamate (blocking lactate dehydrogenase) at Signal 2 could not alter IL-1β secretion or reverse the inhibitory effects of DCA on IL-1β secretion (**Figure 4B**). Blockade of PPP by dehydroepiandrosterone (DHEA, blocking glucose-6-phosphate dehydrogenase) lowered LPS plus ATP-induced IL-1β secretion but could not reverse the effect of DCA on inflammasome activation (**Figure 4C**). These data suggest that neither cytosolic glycolysis nor PPP metabolism is attributable to attenuated ATP-induced IL-1β secretion in PDHK-inhibited macrophages. Blocking mitochondrial pyruvate carrier by UK5099 (**Figure 4D**), PDH-dependent glucose oxidation by CPI-613 (also blocking α-ketogluconate dehydrogenase (α-KGDH) in the TCA cycle) (**Figure 4E**), or inhibition of PC-mediated pyruvate anaplerosis by chlorothricin (**Figure 4F**), suppressed ATP-induced IL-1β secretion, but had no impact on the inhibitory effect of DCA on ATP-induced IL-1β secretion. Inhibiting other metabolic pathways that were altered by PDHK inhibition (**Figure S3A**), for example, the malate-aspartate shuttle by aminooxy acetic acid (AOAA) (**Figure S3B**) or aspartate-argininosuccinate shunt by fumonisin B1 (**Figure S3C**), could not restore IL-1β secretion. These results suggest that PDHK inhibition-mediated NLRP3 inflammasome inactivation does not directly result from enhanced glucose metabolism or the TCA cycle.

**Figure 4.**
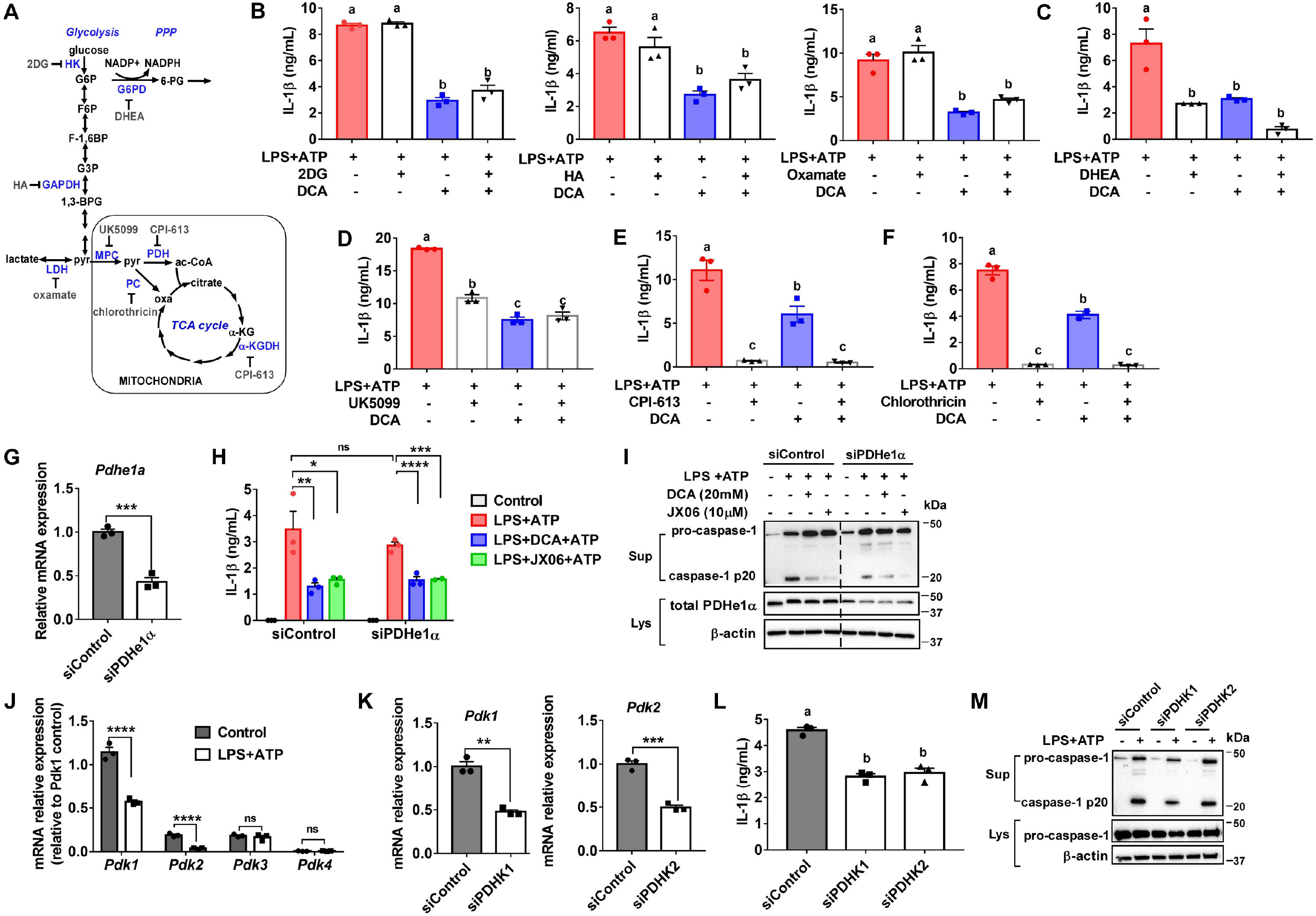
PDHK inhibition-mediated NLRP3 inflammasome inactivation is independent of glucose metabolism or pyruvate dehydrogenase. (**A**) The targeting sites of the pharmacological inhibitors are indicated. (**B-F**) IL-1β secretion from LPS plus ATP-treated BMDMs. The pharmacological inhibitors were added 30 min before ATP stimulation. (**G**) Pdhe1a mRNA expression in control and PDHe1α siRNA transfected elicited peritoneal macrophages (PMs). (**H-I**) IL-1β secretion and caspase-1 cleavage in the culture supernatant from control siRNA and Pdhe1α siRNA transfected PMs treated with or without LPS plus ATP in the presence or absence of DCA or JX06. (**J**) mRNA expression of PDHK isoforms in BMDMs treated with or without LPS plus ATP. Gene expression was normalized to PDHK1 expression in unstimulated macrophages. (**K**) Transcript expression of PDHK isoform 1 and 2 in control and isoform-specific siRNAs (50 µM) transfected PMs, respectively. (**L-M**) IL-1β secretion and caspase-1 cleavage in the culture supernatant from control siRNA and isoform-specific siRNA transfected PMs treated with or without LPS plus ATP. Data are reprehensive of three independent experiments (n=3-4 samples per group). Groups with different letters are significantly different (p<0.05); *P < 0.05; **P < 0.01; ***P < 0.001; ****P < 0.0001; ns: non-significant. One-way ANOVA with post hoc Tukey’s multiple comparisons test (Figures 5B-5F, 5H, 5L); unpaired, two-tailed student’s T-test (Figures 5G, 5J, and 5K).

The initial hypothesis was that PDHK inhibition promotes glucose/pyruvate oxidative metabolism by activating PDC in macrophages, thereby leading to the inactivation of the NLRP3 inflammasome. For this hypothesis to be true, PDH inhibition should reverse the effect of DCA on NLRP3 inflammasome activation. To test this, we knocked down PDHe1α, the PDC subunit that is regulated by PDHK-mediated phosphorylation, using PDHe1α-specific siRNAs. PDHe1α transcript expression showed a 60% reduction in PDHe1α-silenced macrophages (**Figure 4G**). Silencing PDHe1α did not significantly alter the LPS plus ATP-induced IL-1β secretion (**Figure 4H**) but decreased caspase-1 cleavage (**Figure 4I**). Moreover, DCA and JX06 still efficiently lowered LPS plus ATP-induced IL-1β production and caspase-1 cleavage in PDHe1α-silenced macrophages (**Figures 4H-4I**). Together, these data suggest that: a) PDH inhibition cannot reverse PDHK inhibition on NLRP3 inflammasome activity; b) PDHK inhibition-mediated NLRP3 inflammasome inactivation does not result from PDH-induced glucose oxidation. To rule out the off-target effect of the PDHK inhibitors on the NLRP3 inflammasome, we knocked down the predominant macrophage PDHK isoforms to assess whether genetic inhibition of PDHK could mimic the effect of pharmacological inhibitors on NLRP3 inflammasome activity. PDHK1 showed the highest expression in macrophages among the four isoforms, whereas PDHK4 was the lowest abundant isoform at the transcript level (**Figure 4J**). NLRP3 inflammasome stimulation by LPS plus ATP lowered the transcriptional expression of PDHK isoforms 1 and 2, but not 3 or 4 (**Figure 4J**). Based on these results, we silenced the predominant macrophage PDHK isoforms 1 and 2 with isoform-specific siRNAs (**Figure 4K**) and lowered LPS plus ATP-induced IL-1β secretion (**Figure 4L**). Moreover, silencing PDHK isoform 1, and to a much lesser extent, silencing isoform 2, reduced LPS plus ATP-induced caspase-1 cleavage (**Figure 4M)**. Collectively, these data support that PDHK regulates macrophage NLRP3 inflammasome activity by a mechanism independent of PDC-dependent glucose oxidation.

### PDHK inhibition promotes autophagic flux in NLRP3 inflammasome-activated macrophages

Autophagy dampens NLRP3 inflammasome activation partially by clearing assembled inflammasome complexes and removal of damaged mitochondria.^38,39^ We tested whether PDHK inhibition can promote autophagic flux in NLRP3 inflammasome-activated macrophages and, if so, whether autophagy is responsible for the NLRP3 inflammasome inactivation in PDHK-inhibited macrophages. In the absence of bafilomycin (blocking autophagosomal degradation in lysosomes), autophagy-associated protein LC3-II showed less expression in PDHK-inhibited macrophages compared to LPS plus ATP-treated cells. In contrast, in bafilomycin-treated macrophages, DCA or JX06 treatment enhanced the accumulation of both LC3-I and LC3-II (**Figure 5A**). PDHK inhibition also increased expression of the autophagic adaptor protein p62, which facilitates chaperon-mediated autophagy, of mitochondrial Tu translation elongation factor (TUFM), which specifically promotes mitophagy (autophagy of mitochondria) by interacting with Atg5-Atg12, Atg16L1,^40^ or PINK1,^41^ and of Parkin, which mediates mitophagy in a VDAC1 and p62-dependent manner^42^ (**Figure 5B**). These results suggest that PDHK inhibition promotes autophagy and mitophagy in macrophages during inflammasome activation.

**Figure 5.**
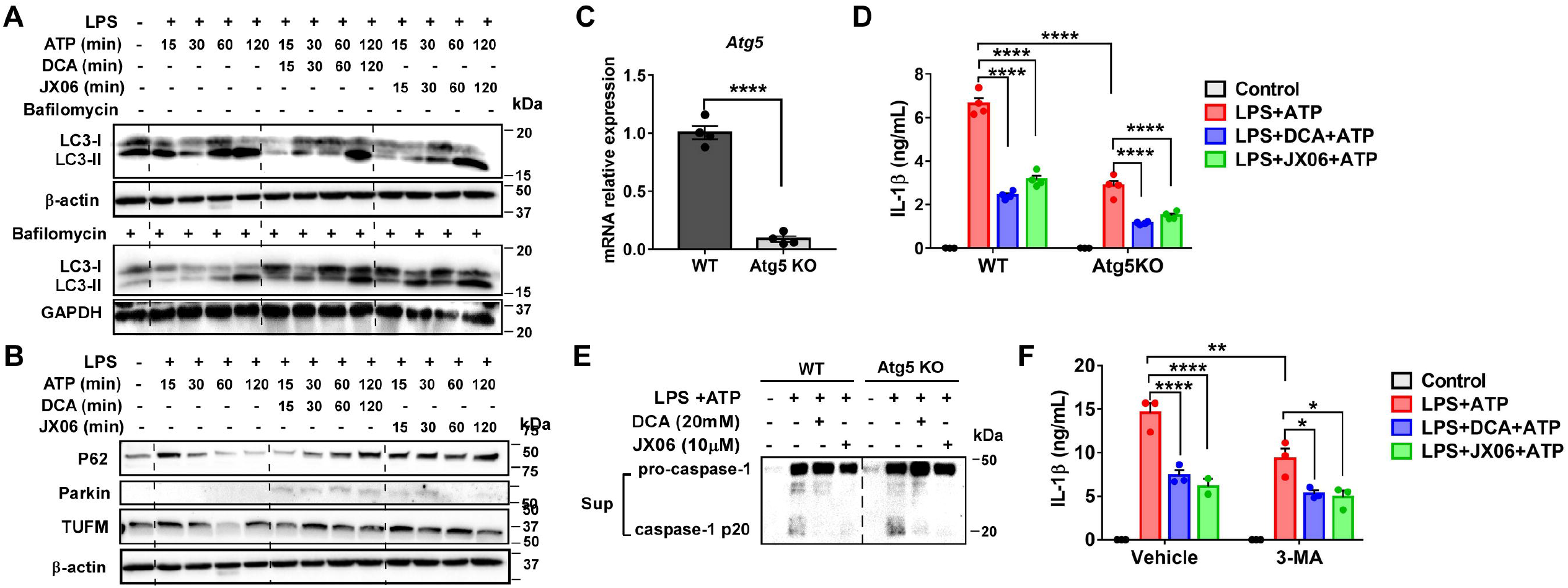
Autophagy is unnecessary for down-regulation of the NLRP3 inflammasome by PDHK inhibition. (**A**) Immunoblotting analysis of LC3 expression in BMDMs treated with or without LPS plus ATP in the presence or absence of DCA or JX06. 50 nM bafilomycin A1 was added together with ATP. (**B**) Immunoblotting analysis of mitophagy marker expression in BMDMs as described in (**A**) except for bafilomycin A1 treatment. (**C**) Atg5 mRNA expression in macrophages. (**D-E**) IL-1β secretion and caspase-1 cleavage in the culture supernatant from WT and Atg5^-/-^ BMDMs. **(F)** 3-methyladenine (3-MA, 5 mM) was added to BMDMs 30 min prior to LPS priming. IL-1β concentration in the culture supernatant was measured by ELISA. Data are reprehensive of three independent experiments (n=3-4 samples per group). *P < 0.05; ****P < 0.0001; ns: non-significant. Unpaired, two-tailed student’s T-test (Figure 5C). Two-way ANOVA with post hoc Tukey’s multiple comparisons test (Figures 5D and 5F).

Next assessed was whether autophagy/mitophagy attenuates NLRP3 inflammasome activation in PDHK-inhibited macrophages. We first used autophagy-deficient macrophages from myeloid cell-specific Atg5 KO mice. Atg5^-/-^ macrophages reduced atg5 transcript expression by 95% relative to control (**Figure 5C**) and had significantly less IL-1β secretion than control (**Figure 5D**), likely due to defective unconventional IL-1β secretion via the secretory autophagy machinery.^43-46^ Interestingly, DCA and JX06 maintained their capacity to reduce IL-1β secretion (**Figure 5D**) and caspase-1 cleavage (**Figure 5E**) in atg5^-/-^ macrophages. Comparable results occurred when 3-methyladenine (3-MA) blocked autophagy (**Figure 5F**). Together, the data suggest that PDHK inhibition promotes autophagy/mitophagy activation independent of decreased NLRP3 inflammasome activity in PDHK-inhibited macrophages.

### PDHK inhibition preserves mitochondrial structure and function in NLRP3 inflammasome-activated macrophages

Mitochondria direct NLRP3 inflammasome activation.^15,17,47,48^ Activating the NLRP3 inflammasome by LPS plus ATP triggered mtROS production (**Figure 6A**), and PDHK inhibition by DCA or JX06 reduced mtROS production by 30% (**Figure 6A**). LPS plus ATP also repressed the methionine-cysteine-glutathione pathways (**Figure S4A**) and decreased NAD^+^ biosynthetic metabolites (**Figure S4B**) that reflect oxidative stress, and JX06 reduced oxidative stress (**Figure S4B**). MitoTracker Deep Red FM mitochondrial imaging using confocal microscopy during NLRP3 inflammasome stimulation converted from a string-like shape to a dot-like shape (**Figure 6B**), as reported.^47,49^ Strikingly, PDHK inhibition by DCA or JX06 maintained mitochondria in a more string-like shape (**Figure 6B**). Further, morphometric analysis of transmission electron microscopy images (**Figure 6C**) indicated reduced mitochondrial length and perimeter in LPS plus ATP-treated vs. control macrophages (**Figures 6D-6E**), suggesting increased mitochondrial fragmentation. PDHK inhibition limited mitochondrial fragmentation, as evidenced by greater mitochondrial length and perimeter in DCA or JX06-treated macrophages (**Figures 6D-6E**). PDHK inhibition did not alter the mitochondrial number, aspect ratio (major axis/minor axis), or roundness compared to NLRP3 inflammasome activation (**Figure S4C**). LPS plus ATP treatment reduced cristae number per mitochondrion (**Figures 6F-6G**) and increased cristae widening (**Figures 6F and 6H**), and PDHK inhibition partially restored cristae density and narrowed cristae (**Figures 6F and 6H**). Collectively, PDHK inhibition improves mitochondrial structure and function and decreases oxidant stress concomitant with reducing NLRP3 activation.

**Figure 6.**
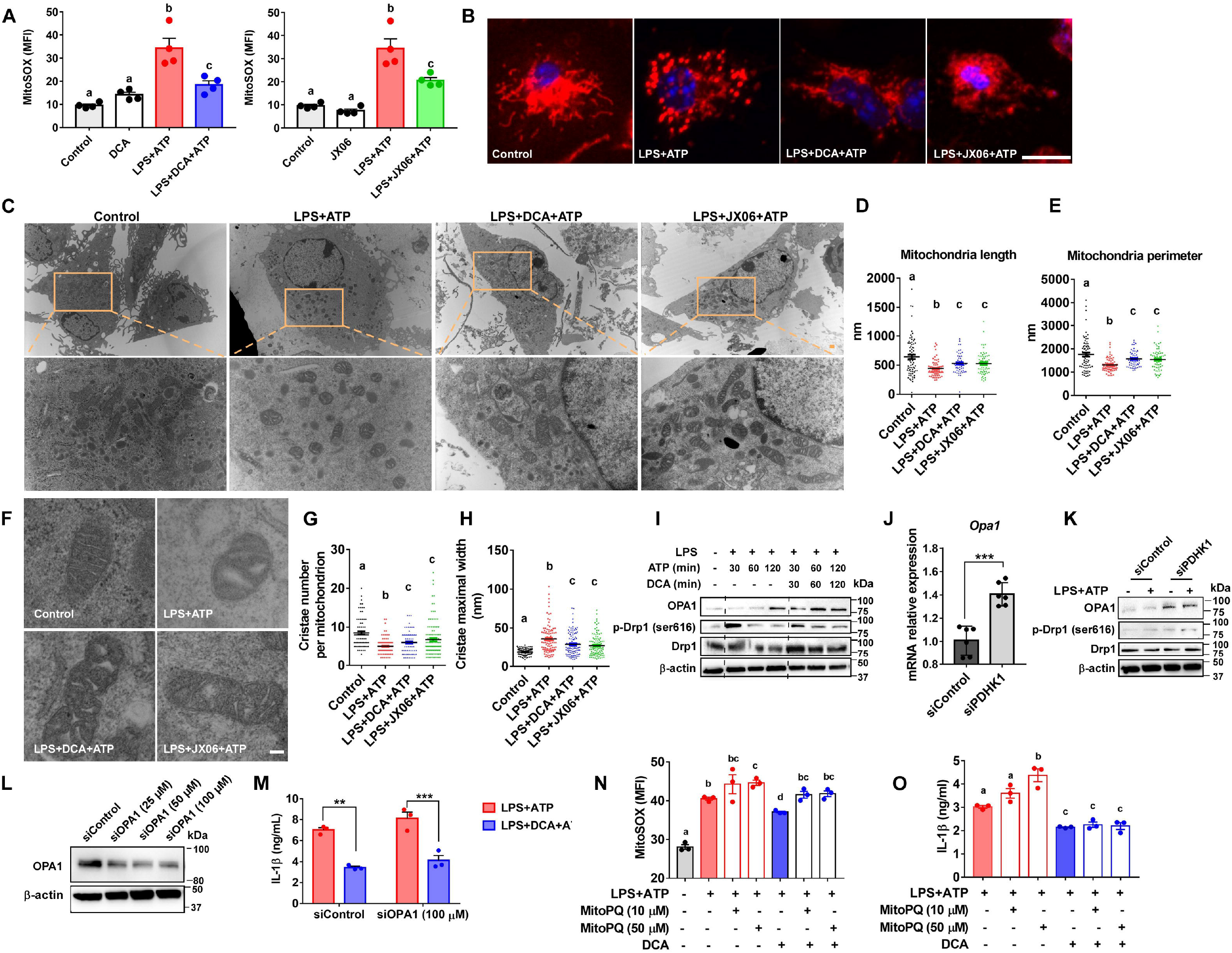
PDHK inhibition-mediated NLRP3 inflammasome inactivation is independent of mitochondrial OPA1 or ROS production. (**A**) BMDMs were stained with MitoSOX and mitochondrial ROS was analyzed by flow cytometry. (**B**) Confocal images of BMDMs stained with MitoTracker Deep Red (red) and DAPI (blue). Scale bar=5 nm. (**C**) Representative transmission electron microscopy (TEM) images of BMDMs. Scale bar=500 nm. (**D-E**) Mitochondrial length and perimeter were quantified. (**F**) Representative mitochondrial TEM images. Scale bar=100 nm. (**G-H**) Mitochondrial cristae number per mitochondrion and cristae width. (**I**) Immunoblotting analysis of p-Drp1 (ser616) and OPA1 in LPS primed BMDMs, followed by ATP plus DCA treatment for 0-120 min. (**J**) Opa1 transcript expression in PDHK1-silenced PMs. (**K**) Immunoblotting analysis of OPA1 and p-Drp1 (ser616) in PMs. PMs were transfected with control siRNA and 50 µM PDHK1 siRNAs before treated with or without LPS plus ATP. (**L**) Immunoblotting analysis of OPA1 protein expression in siRNA transfected PMs. (**M**) IL-1β level in the culture supernatant from siRNAs (100 µM control or OPA1, 72 h) transfected PMs treated with or without LPS plus ATP ± DCA. (**N-O**) MitoParaquat (MitoPQ) were added to LPS primed macrophages 30 min before ATP ± DCA stimulation. BMDMs were stained with MitoSOX (5 µM) and mitochondrial ROS was analyzed by flow cytometry (N). Culture supernatant was analyzed for IL-1β by ELISA (O). Data are reprehensive of three independent experiments (n=3-4 samples per group). ***, P<0.001. Groups with different letters are significantly different (p<0.05) Unpaired, two-tailed student’s T-test (Figure JC). One-way ANOVA (Figures 6A, 6D, 6E, 6G, 6H, 6N, and 6O) or Two-way ANOVA (Figure 6M) with post hoc Tukey’s multiple comparisons test.

Mitochondria fusion and fission are regulated by dynamin-related GTPases, including mitochondrial fission supporting dynamin-related protein 1 (Drp1)^50,51^ and a vital fusion supporting protein optic atrophy protein 1 (OPA1).^52^ Consistent with the increased mitochondrial fragmentation, the protein level of p-Drp1 (Ser616) was markedly increased at 30 min post ATP treatment and its level reduced to the baseline at 60 or 120 min ATP treatment (**Figure 6I**). In contrast, the OPA1 level remained low during 60 min ATP exposure and slightly increased at 120 min ATP stimulation (**Figure 6I**). PDHK inhibition by DCA decreased ATP-induced phospho-Drp1 (Ser616) expression and increased OPA1 protein level (**Figure 6I**). Notably, genetic deletion of PDHK1 also increased OPA1 transcript and protein expression at the basal state and after 1 h ATP stimulation (**Figures 6J-6K**), while p-Drp1 (Ser616) expression did not differ between genotypes, given that its expression declined after 1 h ATP stimulation (**Figure 6K**). Together, PDHK inhibition may promote mitochondrial fusion, preserve cristae, and improve mitochondrial fitness during acute inflammation.

### PDHK inhibition-mediated NLRP3 inflammasome inactivation is independent of mitochondrial fusion or mitochondrial ROS

NLRP3 inflammasome and mitochondrial fission are ERK1/2-dependent,^53^ while mitochondrial fusion prevents NLRP3 inflammasome activation.^54^ To examine the role of mitochondrial fusion on PDHK-regulated inflammasome activity, OPA1 in macrophages was blocked using MYLS22, a selective OPA1 inhibitor, or siRNAs. When added together with ATP and DCA, MYLS22 increased IL-1β secretion (**Figure S4D**), but when added 30 min before LPS priming, MYLS22 could not restore IL-1β secretion (**Figure S4E**). MYLS22 did not affect AIM2 inflammasome activation in the presence or absence of PDHK inhibition (**Figure S4F**). OPA1 siRNAs (**Figure 6L**) could not restore IL-1β secretion in DCA-treated cells (**Figure 6M, Figure S4G**), suggesting that OPA1 protein expression is not required for PDHK inhibition-mediated NLRP3 inflammasome inactivation. Concomitant with rapidly increased Drp1 phosphorylation, LPS plus ATP rapidly activated ERK1/2 at 15 min post ATP stimulation. DCA attenuated ERK1/2 phosphorylation at 15 min (**Figures S4H-S4I**) but inhibiting ERK by U0126 did not affect the inhibitory effect of DCA on IL-1β secretion (**Figure S4J**).

Since PDHK inhibition-mediated NLRP3 inflammasome inactivation might result from attenuated mtROS, mtROS production was induced by mitoParaquat (MitoPQ) at 10 or 50 µM in macrophages treated with or without DCA (**Figure 6N**). MitoPQ at 50 µM increased IL-1β secretion in LPS plus ATP-treated macrophages (**Figure 6O**). Still, DCA treatment lowered IL-1β secretion to the same level in macrophages treated with and without MitoPQ (**Figure 6O**), suggesting that mitochondrial ROS production does not drive PDHK-mediated NLRP3 inflammasome activation.

### PDHK inhibition lowers plasma IL-1β and blood monocyte caspase-1 activity in septic mice

*In vivo* targeting PDHK with DCA, administered as a single dose of 25 mg/kg 24 h post CLP, increases the septic survival rate to 71%, whereas the CLP animals not receiving DCA treatment had a survival rate of only 10% over the 14-day observation period.^33^ Despite the less effect of PDHK inhibition on LPS-induced cytokine expression in BMDMs (**Figures S1H-S1I**), DCA lowered plasma non-inflammasome cytokines, including TNF and IL-6 in CLP mice.^55^ To assess whether DCA promotes sepsis survival by down-regulating the NLRP3 inflammasome during sepsis, we analyzed plasma IL-1β and IL-18 in CLP mice. DCA (25 mg/kg) was given to septic mice 24 h post CLP for 6 h (**Figure 7A**). CLP-induced septic mice showed significantly elevated plasma IL-1β and IL-18 concentrations relative to sham control (**Figures 7B-7C**). NLRP3 inflammasome-induced pyroptosis was assessed by quantifying plasma lactate dehydrogenase (LDH, a cell death marker). At 30 h post-CLP, there was a trend toward increasing plasma LDH in CLP vs. control mice (**Figure 7D**). CLP mice also increased circulating CD115^+^Ly6G^−^ monocytes and CD115^-^Ly6G^+^ neutrophils (**Figures 7E-7G**). A more than 3-fold increase in FLICA (FAM-YVAD-FMK) staining occurred in blood monocytes (CD115^+^Ly6G^-^) and a 2-fold increase in FLICA staining occurred in blood neutrophils (CD115^-^Ly6G^+^) (**Figures 7H-7K**), suggesting elevated caspase-1 cleavage in blood myeloid cells in the CLP vs. sham control mice. DCA treatment significantly decreased plasma IL-1β (**Figure 7B**), lowered plasma LDH (**Figure 7D**), and reduced caspase-1 cleavage in both blood monocytes and neutrophils (**Figures 7H-7K**); DCA also decreased plasma IL-18 slightly (**Figure 7C**). Because IL-1β drives neutrophil recruitment and DCA lowered plasma IL-1β, DCA treatment prevented neutrophilia but not monocytosis in CLP mice (**Figures 7E-7G**). These data suggest that PDHK inhibition attenuates inflammasome-triggered caspase-1 activation and cytokine production during sepsis *in vivo*.

**Figure 7.**
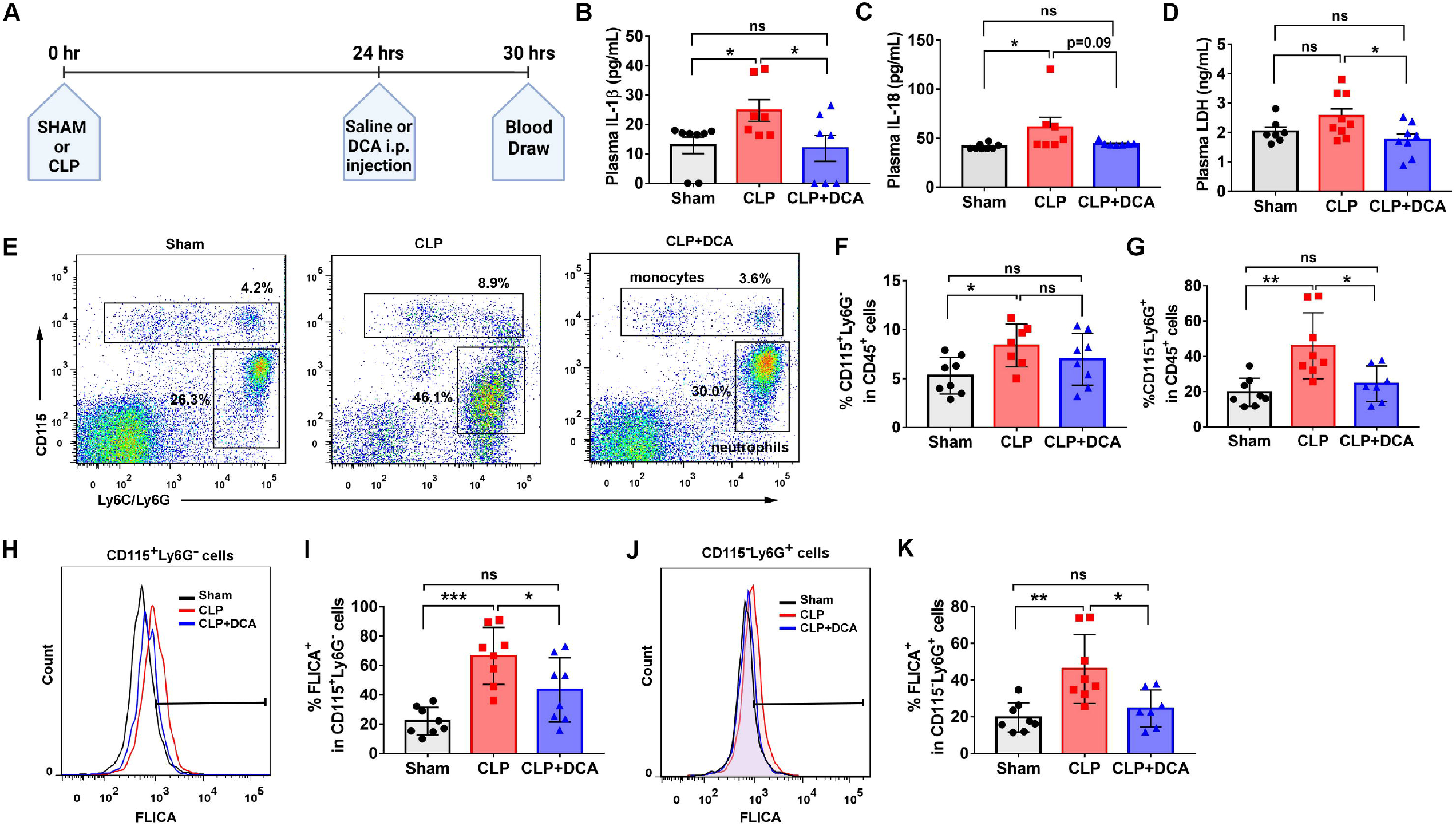
PDHK inhibition lowers plasma IL-1β and blood monocyte and neutrophil caspase-1 activity in septic mice. (**A**) Schematic of mouse treatments. (**B-D**) Plasma IL-1β, IL-18, and lactate dehydrogenase (LDH) were measured by ELISA. (**E-G**) Percentage of blood monocytes (CD115^+^Ly6G^-^) and neutrophils (CD115^-^Ly6G^+^) in CD45^+^ cells were analyzed by flow cytometry. (**H-I**) The percentage of FAM-YVAD-FMK probe (FLICA)^+^ cells in blood monocytes (CD45^+^CD115^+^Ly6G^-^) was analyzed by flow cytometry. (**J-K**) The percentage of FLICA^+^ cells in blood neutrophils (CD45^+^CD115^-^Ly6G^+^) was analyzed by flow cytometry. Each symbol represents an individual mouse (n=7-9 mice). *P < 0.05; **P < 0.01; ***P < 0.001; ns: non-significant, one-way ANOVA with post hoc Tukey’s multiple comparisons test.

## Discussion

Macrophage NLRP3 inflammasome promotes sepsis-associated inflammation^56^ and PDHK inhibits PDC control of glucose oxidation. By phosphorylating the PDHe1α subunit of PDC, PDHK effectively blocks the conversion of pyruvate into acetyl-CoA and limits the TCA cycle function. Pharmacologically targeting PDHK with a pan-PDHK inhibitor DCA enhances septic mouse survival^33^ and rebalances metabolic dysregulation in septic mouse liver^57^ or inflammatory THP1 monocyte sepsis modeling.^58^ Here, our *in vivo* study demonstrates that DCA lowers circulating IL-1β and LDH concentrations, attenuates blood neutrophilia, and reduces blood myeloid cell caspase-1 activity in septic mice, suggesting that PDHK promotes NLRP3 inflammasome activation during sepsis. Our *in vitro* study confirms the role of mitochondrial PDHK in supporting NLRP3 inflammasome activation in murine and human macrophages, occurring independently of PDH-dependent pyruvate oxidation. Moreover, we uncover that PDHK promotes mitochondrial oxidative damage and fragmentation and impairs cristae integrity, supporting NLRP3 inflammasome-induced inflammation and pyroptotic cell death.

In this study, PDHK inhibition markedly reduced NLRP3 inflammasome activation but had a minimal effect on LPS-induced IL-1β, NLRP3, TNF expression, or AIM 2 inflammasome activation, suggesting a pathway-specific effect of PDHK on macrophage inflammation. We initially hypothesized that PDHK supports NLRP3 inflammasome activation by downregulating PDH-dependent pyruvate metabolism. Interestingly, blockade of pyruvate transport into mitochondria, inhibition of PC-dependent pyruvate anaplerosis, or inhibition of PDH-dependent pyruvate oxidation at Signal 2 all markedly reduced IL-1β secretion. Consistent with our findings, mitochondrial pyruvate-derived citrate transports into the cytosol for fatty acid synthesis to support the NLRP3 inflammasome activation.^59^ PDH-dependent pyruvate flux enhances LPS-induced macrophage inflammation.^60^ Thus, our results and other published work suggest that mitochondrial pyruvate metabolism supports NLRP3 inflammasome activation. Notably, inhibition of mitochondrial pyruvate metabolism could not reverse the suppressive effect of DCA on inflammasome activation. Moreover, both DCA and JX06 effectively lowered IL-1β secretion in PDHe1α-silenced macrophages. Our findings suggest a role for PDHK in regulating the NLRP3 inflammasome, independent of the phosphorylation of PDH for pyruvate oxidation.

Our results clearly showed that PDHK inhibition protects macrophages against NLRP3 inflammasome activation-induced mitochondrial oxidative damage and dysfunction, as evidenced by 1) reduced mitochondrial fragmentation, 2) preserved cristae ultrastructure, 3) attenuated mtROS production, and 4) enhanced ATP generation. Intriguingly, a recent study reported that mitochondrial electron transport chain (ETC)-generated ATP, but not ROS, is required for activating NLRP3 inflammasome.^26^ In our study, down-regulation of mtROS is not necessary for PDHK inhibition-mediated NLRP3 inflammasome inactivation, given that DCA can equally lower LPS plus ATP-induced IL-1β secretion in macrophages treated with or without mitoPQ. Moreover, because PDHK inhibition enhances glycolysis, mitochondrial respiration, and total ATP production in NLRP3-activated macrophages, it is unlikely that the down-regulation of NLRP3 inflammasome activation in PDHK-inhibited cells results from decreased mitochondrial TCA and ETC activities. Controversial results exist regarding the role of mitochondrial dynamics in macrophage NLRP3 inflammasome activation.^53,54,61^ In our study, PDHK-inhibited macrophages decreased mitochondrial fission protein p-Drp1 and conversely increased mitochondrial fusion protein OPA1 expression, in parallel with the improved mitochondrial morphology and cristae ultrastructure during NLRP3 inflammasome activation. However, knockdown of OPA1 or indirect inhibition of Drp1-mediated mitochondrial fission by blockade of ERK1/2 signaling pathway could not reverse the effect of DCA on NLRP3 inflammasome activation, suggesting that PDHK inhibition-mediated NLRP3 inflammasome inactivation is independent of OPA1 expression or mitochondrial elongation.

A generalized metabolic defect occurs in leukocytes at the later stage of human sepsis.^62^ Interestingly, LPS-primed macrophages elevated Warburg glycolysis without altering mitochondrial respiration, mimicking the initial step of sepsis.^62^ LPS plus ATP-induced NLRP3 inflammasome activation parallels broad metabolic defects in macrophages, mimicking the metabolic paralysis that rapidly emerges and persists during sepsis. PDHK inhibition at Signal 2 broadly rewired cellular metabolic pathways and rescued metabolic defects by increasing glucose uptake and flux through glycolysis and TCA cycle via both PDC and PC anaplerosis portals for energy production and cell survival. Mitochondrial cristae shape regulates respiratory chain super-complex stability and assembly.^63^ Although mitochondrial dynamics has no impact on PDHK-regulated NLRP3 inflammasome activity, our results suggest a role for PDHK in supporting mitochondrial oxidative stress, mitochondrial fission over fusion, and cristae remodeling upon NLRP3 inflammasome stimulation. The underlying mechanisms of mitochondrial structural retrieval by PDHK inhibition require further investigation. As a potential guide, PDHK4 can sequester NF-κB/p65 in the cytoplasm via a direct protein-protein interaction in hepatocytes.^64^ PDHK4 also promotes mitochondrial fission via a PDHK4-SEPT2-DRP1 axis.^65^ These studies indicate the dual location and function of PDHK4. PDHK1 is the most abundant isoform expressed in macrophages. Inhibition of PDHK by DCA or genetic deletion up-regulated OPA1 expression at both transcript and protein levels, making it likely that PDHK1 translocates to the cytoplasm to regulate mitochondrial fission/fusion protein expression by phosphorylating transcriptional factors or signaling molecules involved in this process.

Autophagy regulates organelle and protein turnover by both degradative and secretory mechanisms. Degradative autophagy inhibits NLRP3 inflammasome activation by limiting mitochondrial DNA release,^17^ inducing inflammasome complex degradation,^66,67^ or clearing damaged mitochondria.^15^ In contrast, secretory autophagy facilitates the extracellular delivery of IL-1β through an unconventional secretory pathway.^43-46^ Therefore, in the autophagy-deficient macrophages, the level of extracellular IL-1β is the net result of the increased IL-1β cleavage due to enhanced inflammasome activation and the attenuated IL-1β secretion due to the impaired unconventional secretory pathway. Nonetheless, our results showed that PDHK inhibition promotes autophagic flux and mitophagy in NLRP3 inflammasome-stimulated macrophages. Despite the indispensable role of autophagy in attenuating IL-1β secretion by PDHK inhibition, the enhanced autophagic flux, especially mitophagy, paralleled improved mitochondrial morphology and function in PDHK-inhibited macrophages, suggesting that PDHK inhibition, in part, promotes mitochondrial homeostasis by activating mitophagy.

In conclusion, we discover non-canonical roles for mitochondrial PDHK in supporting mitochondrial oxidative damage and NLRP3 inflammasome activation independent of its canonical regulation of limiting pyruvate oxidation. Although the detailed mechanisms need further elucidation, targeting PDHK dampens NLRP3 inflammasome activity, reverses metabolic paralysis, and maintains mitochondrial fitness in inflamed macrophages. These aspects promote macrophage homeostasis and health in sepsis, making PDHK inhibition by DCA/JX06 translationally valuable. In addition to sepsis, PDHK has emerged as a critical target in other disease states, including diabetes,^68^ cardiovascular diseases,^69^ and aging.^70^ Hence, our study provides evidence for the therapeutic targeting of PDHK to treat many inflammatory conditions.

### Limitations of the study

In this study, we utilized various pharmacological inhibitors to examine the effect of the metabolic pathways on NLRP3 inflammasome activation. There is potential that non-specific untargeted effects exist and interfere with the data interpretation. Also, we blocked each targeted pathway using a pathway-specific inhibitor when we examined the impact of metabolic shift induced by PDHK inhibition on the NLRP3 inflammasome. We do not rule out that there could be a combinatory effect through multiple pathways on PDHK-regulated NLRP3 inflammatory activation.

## Acknowledgments

The authors gratefully acknowledge Ken Grant, Paula Graham, and Debbie Golden of the Cellular Imaging Shared Resource of Atrium Health Wake Forest Baptist for technical assistance with confocal and transmission electron microscopy. The authors would like to acknowledge pilot funding support from the Center for Redox Biology and Medicine and the Cardiovascular Sciences Center at WFSM. This study was supported by NIH R35 GM126922 (CEM), NIH R01 CA248037 (HKL), NIH R01 HL132035 (XZ), and the National Center for Advancing Translational Sciences of NIH under UL1TR001420 (Research Assistant fund to XZ). AKM was supported by NIH T32GM127261 and NIH T32 AI007401 training grants.

## Author Contributions

X.Z. conceived the study; A.K.M., Z.W., W.H., Q.Zhao, M.Z., Q.Zhang, and F.L. conducted the experiments; L.D. conducted the ^13^C glucose flux assay and edited the manuscript; J.L. conducted the high-resolution metabolomics analysis and edited the manuscript; R.K.M provided advice related to TEM; J.W.L. provided advice related to metabolomics analysis and helped with data discussion; M.A.Q, Q.S., D.F., H.K.L., C.M.F., and C.E.M helped with data discussion and provided comments and revisions; Z.W. wrote the methods and edited the manuscript; A.K.M. and X.Z. drafted the manuscript; X.Z. wrote and finalized the manuscript.

## Inclusion and diversity statement

We support inclusive, diverse, and equitable conduct of research.

## Star Methods

### RESOURCE AVAILABILITY

#### Lead contract

Further information and request for resources and reagents should be directed to and will be fulfilled by the lead contact, Xuewei Zhu.

#### Materials availability

All unique reagents generated in this study are available from the lead contact without restriction.

#### Data and code availability

- Targeted metabolomics, untargeted metabolomics, and 13C-glucose tracing data are available at the NIH Common Fund’s National Metabolomics Data Repository (NMDR) website, the Metabolomics Workbench,^71^ https://www.metabolomicsworkbench.org, where it has been assigned Project ID PR001524. The data can be accessed directly via it’s Project DOI: 10.21228/M8Q13W.
- This paper does not report original code.
- Any additional information required to reanalyze the data reported in this paper is available from the lead contact upon request.

### EXPERIMENTAL MODEL AND SUBJECT DETAILS

#### Mice

C57BL/6J (stock 000664), LysMcre (stock 004781), and Gsdmd^-/-^ (stock 032410) mice were purchased from Jackson Laboratories. Atg5^flox/flox^ mice were purchased from Riken BioResource Center (stock RBRC02975). Heterozygous atg5 macrophage-specific knockout (atg5^flox/+^Lyscre^+^) mice were generated by crossing atg5^flox/flox^ mice and LysMcre mice. Homozygous atg5 macrophage-specific knockout (atg5^flox/flox^LysMcre^+^) and WT control mice (atg5^flox/flox^) were generated by intercrossing the heterozygous mice, as described previously.^72^ Mice were housed in a pathogen-free facility on a 12 h light/dark cycle and received a standard laboratory diet. Cecal ligation and puncture (CLP) were performed to induce intra-abdominal polymicrobial infection in C57BL/6J male mice at the age of 12 weeks as described previously.^33^ Sham-operated mice were used as a control. DCA (25 mg/kg) or saline was injected intraperitoneally (i.p.) 24 h post-CLP. Mice were euthanized 30 hours post CLP. All animal experimental protocols were approved by the Wake Forest University Animal Care and Use Committee.

#### Cells

Bone marrow-derived macrophages (BMDM) and thioglycollate-elicited peritoneal macrophages were cultured as described previously.^73,74^ Briefly, mouse bone marrow was isolated from C57BL/6J mice (male or female, 10-20 weeks old), atg5^flox/flox^ and atg5^flox/+^Lyscre^+^ (male, 12 weeks old), or WT and Gsdmd^-/-^ (male, 10 weeks old) mice. Bone marrow cells were cultured in low glucose DMEM supplemented with 30% L929 cell-conditioned medium, 20% FBS, 2 mM L-glutamine, 1 mM sodium pyruvate, 100 U/ml penicillin, and 100 µg/ml streptomycin for 6-7 days until the cells reached confluence. BMDMs were then reseeded in culture dishes overnight in RPMI-1640 medium containing 1% Nutridoma-SP medium (Sigma-Aldrich) before any treatment. Thioglycollate-elicited peritoneal macrophages (PMs) were harvested from C57BL/6J mice (male or female, 10-15 weeks old) by washing peritoneal cavities using cold PBS 3 days after i.p. injection of 1 ml of 10% thioglycollate (Sigma-Aldrich). After a 2 h Incubation in RPMI-1640 media containing 10% FBS, 100 U/ml penicillin, and 100 µg/ml streptomycin, floating cells were removed, and adherent macrophages were used for experiments. Peripheral blood mononuclear cells (PBMCs) were isolated from fresh blood from a male healthy donor using 1:1 ratio of blood to Histopaque 1077. PBMCs were subsequently cultured in RPMI-1640 medium supplemented with M-CSF (100 ng/mL), 20% FBS, 2 mM L-glutamine, 1 mM sodium pyruvate, 100 U/ml penicillin, and 100 µg/ml streptomycin for 6-7 days until the cells reach confluence. PBMCs-derived macrophages were replaced in culture dishes before inflammatory stimulation. The study protocol was approved by the Wake Forest University Institutional Review Board.

### METHOD DETAILS

#### Cell treatment

To induce NLRP3 inflammasome activation, macrophages were first primed with 300 ng/ml LPS (E. coli 0111; B4, Sigma-Aldrich) before stimulated with 5 mM ATP (Sigma-Aldrich) for 15 min to 120 min, 10 µM nigericin (InvivoGen) for 1 h, 0.6 mg/ml monosodium urate crystals (MSU) (InvivoGen) for 6 h in RPMI-160 medium, as indicated in the figure legends. To assess the AIM2 inflammasome, BMDMs were first primed with 300 ng/ml LPS for 3 h before stimulated with poly (dA:dT) (2.5 µg/mL, InvivioGen) complexed with transfection reagent LyoVec for 7 h. To inhibit PDHK, macrophages were pretreated for 30 min before LPS priming or after LPS priming (together with inflammasome stimuli as described above) with DCA (5-30 mM, Sigma-Aldrich), JX06 (5-40 µM, Rocris) for an indicated time as described in the figure legends. In some experiments, LPS-primed macrophages were treated with AOAA (1 mM, Cayman Chemical), bafilomycin A1 (50 nM, In vivoGen), chlorothricin (100 µM, Cayman Chemical), CPI-613 (200 µM, Cayman Chemical), 2-DG (10 mM, Sigma-Aldrich), DHEA (200 µM, Sigma-Aldrich), etomoxir (50 µM, Cayman Chemical), fumonisin B1 (15 µM, Cayman Chemical), glycine (5 mM, Sigma-Aldrich), heptelidic acid (15 µM, Cayman Chemical), 3-MA (5 mM, Sigma-Aldrich), ML385 (5 µM, MedChemExpress), SC-26196 (10 µM, Cayman Chemical), sodium oxamate (40 mM, Santa Cruz Biotechnology), U0126 (25 µM, Calbiochem), or UK5099 (5 µM, Cayman Chemical), as indicated in the figure legends. PBS, DMSO, or ethanol was used as vehicle control in the inhibitor experiments.

#### Flow cytometry

After removing the red blood cells using ACK lysing buffer (Gibco), peripheral white blood cells were stained with FLICA 660-YVAD-FMK (1:150 dilution, ImmunoChemistry Technologies), CD115-PE (eBioscience, no. 12-1152), Gr1(Ly6C/G)-PerCP-Cy5.5 (BioLegend, no. 108427), and CD45-APC (BD Pharmingen, no. 559864). Data were acquired on a BD FACS Canto II instrument (BD Biosciences) and analyzed using FlowJo analytical software (version 7.6.5, TreeStar). To measure cell death, BMDMs were stained with 7-AAD (Invitrogen, cat# V35123) or PI (Invitrogen, Cat# P3566) according to the manufacturer’s instructions. To quantify glucose uptake, BMDMs were exposed to 100 µg/ml 2-NBDG (Cayman Chemical, Cas#186689-07-6) after LPS priming for 30 min. Data were acquired on a BD FACSCalibur and analyzed using FlowJo analytical software (version 7.6.5, TreeStar).

#### Metabolic analysis

##### High-resolution untargeted metabolomics

LPS (300 ng/ml, 3 h)-primed BMDMs were stimulated with 5 mM ATP in the presence or absence of 10 µM JX06 for 45 min. Unstimulated macrophages were used as control. Macrophages were lysed, and polar metabolites were extracted using methanol and H_2_O (80:20; HPLC Grade; Sigma-Aldrich).^75^ Extracts were dried in a vacuum concentrator at room temperature, followed by liquid chromatography using Ultimate 3000 UHPLC (Dionex) coupled to high-resolution-mass spectrometry for metabolite profiling using the Q Exactive Plus-Mass spectrometer (QE-MS, Thermo Scientific).^76^ Commercial software Sieve 2.2 (Thermo Scientific) was using for peak extraction and integration. Liquid chromatography-mass spectrometry peak areas were normalized to sample protein mass measured by the BCA assay. Pathway analysis of metabolites was conducted using the MetaboAnalyst v.5.0.

##### Targeted metabolomics

LPS (300 ng/ml, 3 h)-primed BMDMs were stimulated with 5 mM ATP in the presence or absence of 10 µM JX06 for 30 min. Unstimulated macrophages were used as control. Macrophages were lysed, and polar metabolites were extracted using methanol and H_2_O (80:20; HPLC Grade; Sigma-Aldrich).^75^ 500 µl of cell extracts and 20 µl of MES (Thermo Fisher Scientific) internal standard solution (10 ng/µl) were mixed and dried under vacuum and reconstituted for analysis in 100 µl of ultrapure water (Optima, ThermoFisher Scientific). The analysis was performed on a Shimadzu Nexera UHPLC system coupled with a Shimadzu LCMS-8050 triple-quadrupole mass spectrometer (Kyoto, Japan). Two LC-MS/MS methods (Ion-paring and PFPP) were employed to measure the targets. Data were normalized to sample protein mass measured by the BCA assay.

##### In vitro ^13^C glucose tracing

LPS (300 ng/ml, 3 h)-primed BMDMs were stimulated with or without 5 mM ATP in the presence or absence of 10 µM JX06 for 30 min. ^13^C tracing started by replacing medium with glucose-free medium supplemented with 2.06 g/L [U-^13^C]-glucose (Cayman Chemical) for 90 min. Cells were washed with ice-cold saline and intracellular metabolites were extracted using cold methanol and H2O (80:20; HPLC Grade; Sigma-Aldrich) as described above.^75^ The samples were pre-dissolved into 40 µL solution (LC-water, methanol, Acetonitrile 2:1:1) before injection. For each sample, 3 µL solution was injected into the LC-MS for analysis. The HPLC analysis of the isotope-labeled samples was performed using Ultimate 3000 UHPLC (Dionex) as described.^77^ The mass spectrometry analysis was performed using Q Exactive Plus mass spectrometer (Thermo Fisher Scientific). The mass spectrometers were equipped with a HESI probe and operated in the positive/negative switching mode. When Q Exactive Plus mass spectrometer was used, the relevant parameters were as listed: heater temperature, 120 °C; sheath gas, 30; auxiliary gas, 10; sweep gas, 3; spray voltage, 3.0 kV; capillary temperature, 320°C; S-lens, 55. The resolution was set at 70,000 (at m/z 200). Maximum injection time (max IT) was set at 200 ms and automated gain control (AGC) was set at 3 × 10^6^. The LC-MS peak extraction and integration of the raw data were performed using commercially available software Sieve 2.0 (Thermo Fisher Scientific). The integrated peak area was used to calculate ^13^C enrichment. Natural abundance correction was performed using software R with Bioconductor R package IsoCorrectoR.

##### Seahorse assays

2 × 10^5^ BMDMs were plated into each well of Seahorse XF96 cell culture microplates (Agilent Technologies) and cultured overnight in in 1% Nutridoma SP medium before treated with or without 300 ng/ml LPS for 3 h plus 5 mM ATP in the presence or absence of 20 mM DCA or 10 µM JX06 for 30 min. Cells were washed with designed seahorse media and continued to be incubated in the seahorse media for another 30 min. Extracellular acidification rate (ECAR), oxygen consumption rate (OCR), and real-time ATP production rate in BMDMs were measured by using glycolysis stress kit, mitochondrial stress kit, and real-time ATP rate assay kit, respectively, with a Seahorse XF96 Extracellular Flux Analyzer (Agilent Technologies). 10 mM glucose, 1 µM oligomycin, 1.5 µM fluoro-carbonyl cyanide phenylhydrazone (FCCP), 100 nM rotenone plus 1 µM antimycin A, or 50 mM 2DG (all the compounds were from Agilent Technologies) were sequentially injected into the microplates according to the manufacturer’s instructions. After the assay, 3 µM Hoechst (Life Technologies) was added to each well to stain nuclei for cell counting. Results were collected with Wave software version 2.6 (Agilent Technologies). Data were normalized to cell numbers.

#### Microscopy analysis

##### DIC imaging

DIC images of BMDMs were obtained using a Zeiss Axio Vert.A1 microscope with AxioCam ICc3 camera. Cells in more than 5 fields of a 20x objective were analyzed for cell size using NIH ImageJ software (NIH).

##### Confocal imaging

BMDMs seeded in a glass chamber slide were first primed with LPS (300 ng/ml or 3 h. BMDMs were then stimulated with or without 5 mM ATP in the presence or absence of 10 µM JX06 for 45 min before stained with 100 nM MitoTracker Deep Red (Invitrogen) for 15 min at 37 °C in dark. Macrophages were then washed and fixed with 4% paraformaldehyde. Nuclei were stained using DAPI. Confocal images were obtained with a FV1200 laser-scanning confocal microscope (Olympus).

##### Transmission electron microscopy (TEM) imaging

LPS (300 ng/ml, 3 h)-primed BMDMs were stimulated with or without 5 mM ATP in the presence or absence of 10 µM JX06 for 45 min. BMDMs were then subjected to TEM imaging in the Wake Forest Baptist Health electron microscope core Lab. Briefly, macrophages were fixed in 2.5% glutaraldehyde in 0.1 M Millonig’s phosphate buffer pH 7.3 for a minimum of 1 h, wash with buffer, and post-fixed in % osmium tetroxide for 1 h. After dehydration, resin infiltration, and cure in a 70 °C oven overnight, cells were sectioned on a Reichert-Jung Ultracut E ultramicrotome. The sections were then stained with lead citrate and uranyl acetate and viewed with an FEI. Tecnai Spirit TEM was operating at 80 kV. Images were obtained with an AMT 2Vu CCD Camera. Mitochondrial morphology was analyzed on images with 4800x or 18500x magnification. More than 20 cells or 70 to 100 mitochondria per group were analyzed to quantify mitochondrial or cristae ultrastructure changes using NIH ImageJ software (NIH).

#### Mitochondrial ROS (mtROS) measurement

mtROS production in BMDMs was measured by MitoSOX (Fisher Scientific, no. M36008) staining (5 µM for 15 min at 37°C). Macrophages were washed with PBS and scraped off the dishes. Data were acquired on a BD FACSCalibur and analyzed using FlowJo analytical software (version 7.6.5, TreeStar).

#### siRNA transfection

25-100 nM control, PDHK1, PDHK2, PDHK4, PDHE1a, or OPA1 ON-Targetplus siRNAs (Dharmacon, Lafayette, CO) were transfected into elicited peritoneal macrophages isolated from WT C57BL/6 mice with DharmaFECT 1 transfection reagent (Dharmacon, Lafayette, CO) according to the manufacturer’s protocol. siRNA transfection efficiency was quantified by qPCR after 48 h transfection or western blotting after 72 h transfection.

#### ELISA

Mouse IL-1β, IL-18, TNF, IL-6, or LDH were measured in macrophage culture supernatants or mouse plasma using commercially available ELISA kits according to the manufacturer’s instructions.

#### qPCR

Total RNA was isolated from macrophages using Trizol reagent (Thermo Fisher Scientific). cDNA was synthesized with Omniscript RT Kit (Qiagen). The relative mRNA expression level of each target gene was quantified by qPCR using KAPA SYBR fast qPCR master mix (Roche). Data were normalized to GAPDH and expressed relative to control unstimulated macrophages. The sequences of the primers are listed in Table S1.

#### Western Blotting

Macrophage cell lysates were lysed in radioimmunoprecipitation assay (RIPA) buffer containing protease inhibitor cocktails and PhosSTOP (Roche). Protein concentration was measured using the BCA Protein Assay Kit (Pierce). Primary antibodies used were anti-caspase-1 p20 (AdipoGen, no. AG-20B-0042; 1:1000), LC-3 (Novus Biologicals, no. NB100-2220; 1:500), P62 (Novus Biologicals, no. NBP1-48320; 1:4000), NLRP3 (Adipogen, no. AG-20B-0014; 1:1000), ASC (Adipogen, no. AG-25B-0006; 1:1000), PDHe1α (ThermoFisher, no. PA5-21536; 1:1000), TUFM (Novus Biologicals, no. CL2242; 1:500), GAPDH (Invitrogen, no. MA5-15738; 1:10,000), OPA1 (Cell Signaling, no. CST80471; 1:1000), p-DRP1 (Ser616) (Cell Signaling, no. 3455S; 1:500), p-TBK1 (Ser172) (Cell Signaling, no. CST5483; 1:1000), p-Erk1/2 (Cell Signaling, no. 9101; 1:1000), Erk1/2 (Cell Signaling, no. 9102; 1:1000), and β-actin (Sigma-Aldrich, no. A5441; 1:5000). Immunoblots were visualized with the Supersignal substrate system (Thermo Fisher Scientific), and chemiluminescence was captured using the ChemiDox MP imaging system (Bio-Rad) or an LSA-3000 imaging system (Fujifilm Life Science).

### QUANTIFICATION AND STATISTICAL ANALYSIS

In vitro experiments were performed in triplicate or quadruplicate except for the seahorse assays which were performed in at least quadruplicate. Individual dots in each graph represented individual data points for *in vivo* and *in vitro* experiments. At least two independent experiments were performed for each assay except for the metabolomics analysis. Statistical analysis was performed using GraphPad Prism software 7 (GraphPad Software). Data are presented as the mean ± SEM unless indicated otherwise. Differences were compared with Student’s t-test, one-way ANOVA, or two-way ANOVA with post hoc Tukey’s multiple comparison test. Significant differences were regarded as p□<□0.05, p□<□0.01, p□<□0.001, or p<0.0001, indicated in the figure legends.

