## Supplemental information for "Pyruvate dehydrogenase kinase supports macrophage NLRP3 inflammasome activation during acute inflammation"

**Table SI. Forward and reverse primers used in qPCR, Related to Figures 1, 4-6**

| Gene name | Forward primer (5'→3') | Reverse primer (5'→3') |
|---|---|---|
| Gapdh | TGTGTCCGTCGTGGATCTGA | CCTGCTTCACCACCTTCTTGAT |
| Nlrp3 | TCCCAGACACTCATGTTGCC | GTCCAGTTCAGTGAGGCTCC |
| Il1b | GTCACAAGAAACCATGGCACAT | GCCCATCAGAGGCAAGGA |
| Atg5 | TGTGCTTCGAGATGTGTGGTT | GTCAAATAGCTGACTCTTGGCAA |
| Pdhe1a | TGTGACCTTCATCGGCTAGAA | TGATCCGCCTTTAGCTCCATC |
| Pdk1 | GGACTTCGGGTCAGTGAATGC | TCCTGAGAAGATTGTCGGGGA |
| Pdk2 | AGGGGCACCCAAGTACATC | TGCCGGAGGAAAGTGAATGAC |
| Pdk3 | TCCTGGACTTCGGAAGGGATA | GAAGGGCGGTTCAACAAGTTA |
| Pdk4 | CCATGAGAAGAGCCCAGAAGA | GAACTTTGACCAGCGTGTCTACAA |
| Opa1 | TGGAAAATGGTTCGAGAGTCAG | CATTCCGTCTCTAGGTTAAAGCG |

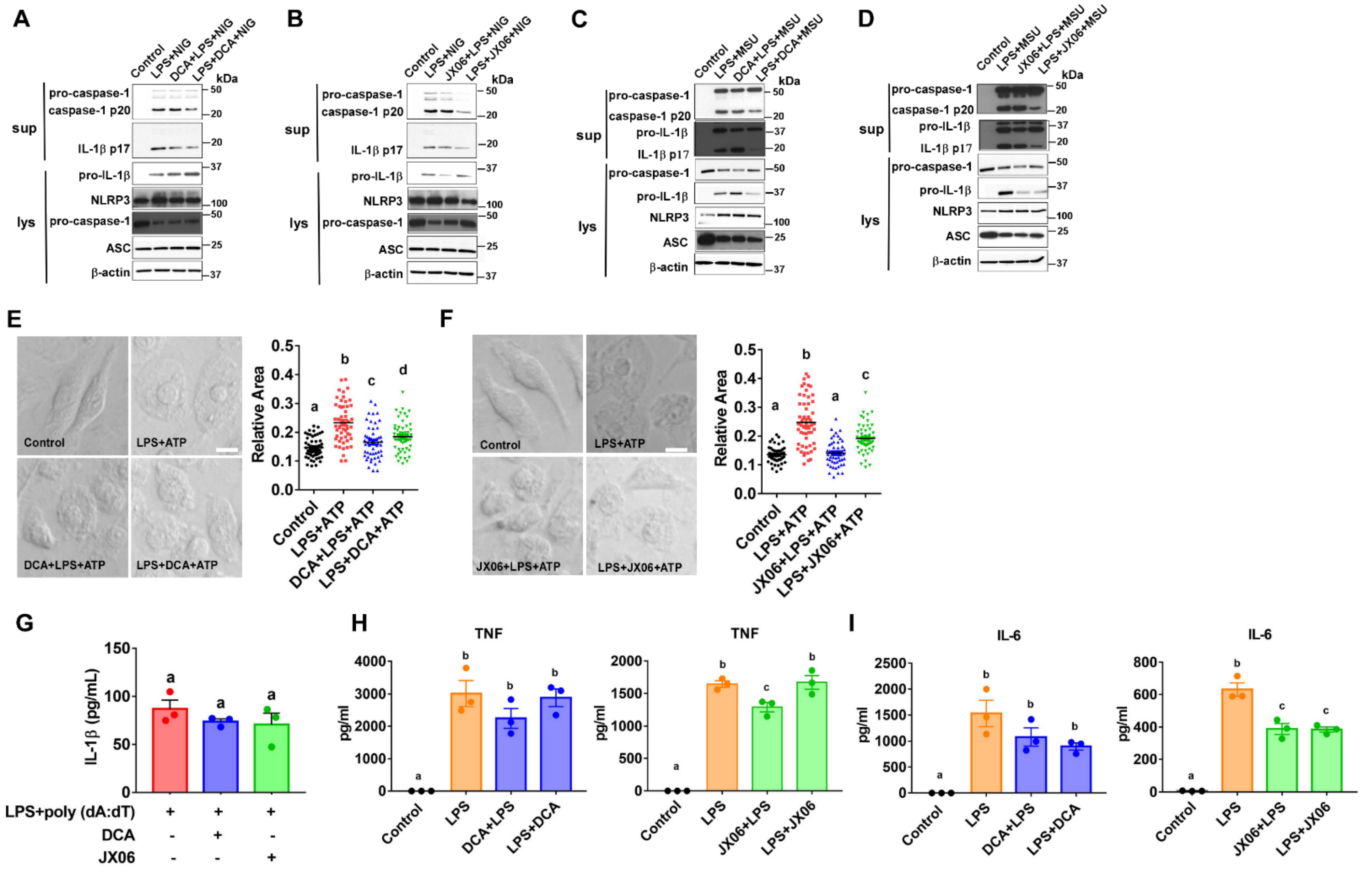

**Figure S1. PDHK inhibition has a more profound inhibitory effect on the NLRP3 inflammasome, Related to Figure 1.**

(**A-D**) BMDMs were first primed with 300 ng/ml LPS for 3 h before stimulated with or without 10 μM nigericin (NIG) for 1 h or 0.6 mg/ml monosodium urate crystals (MSU) for 6 h, respectively. 20 mM DCA, 10 μM JX06, or DMSO (vehicle control for JX06) was added before or after LPS priming, together with the NLRP3 inflammasome stimulus. Protein expression in the culture supernatant (Sup) and whole-cell lysates (Lys) was analyzed by immunoblotting.

(**E-F**) BMDMs were first primed with 300 ng/ml LPS for 3 h before stimulated with or without 5 mM ATP for 45 min. 20 mM DCA, 10 μM JX06, or DMSO (vehicle control for JX06) was added before or after LPS priming, together with the NLRP3 inflammasome stimulus. Phase-contrast images of macrophages were captured by a microcopy, and the relative area of macrophages was analyzed. Scale bar=10 μm.

(**G**) BMDMs were first primed with 300 ng/ml LPS for 3 h before stimulated with or without 2.5 μg/ml poly (dA:dT)/LyoVec for 7 h. 20 mM DCA, 10 μM JX06, or DMSO (vehicle control for JX06) was added together with poly (dA:dT). IL-β concentrations in the culture supernatant were analyzed by ELISA.

(**H-I**) Cytokine secretion from BMDMs treated with or without 300 ng/ml LPS for 4 h. 20 mM DCA or 10 μM JX06 were added 30 min before LPS (DCA+LPS or JX06+LPS) or 3 h after LPS stimulation (LPS+DCA or LPS+JX06). Cytokine concentrations in the culture supernatant were measured by ELISA.

Data are reprehensive of three independent experiments (n=3-4 samples per group). Groups with different letters are significantly different ($p<0.05$); One-way ANOVA with post hoc Tukey's multiple comparisons test.

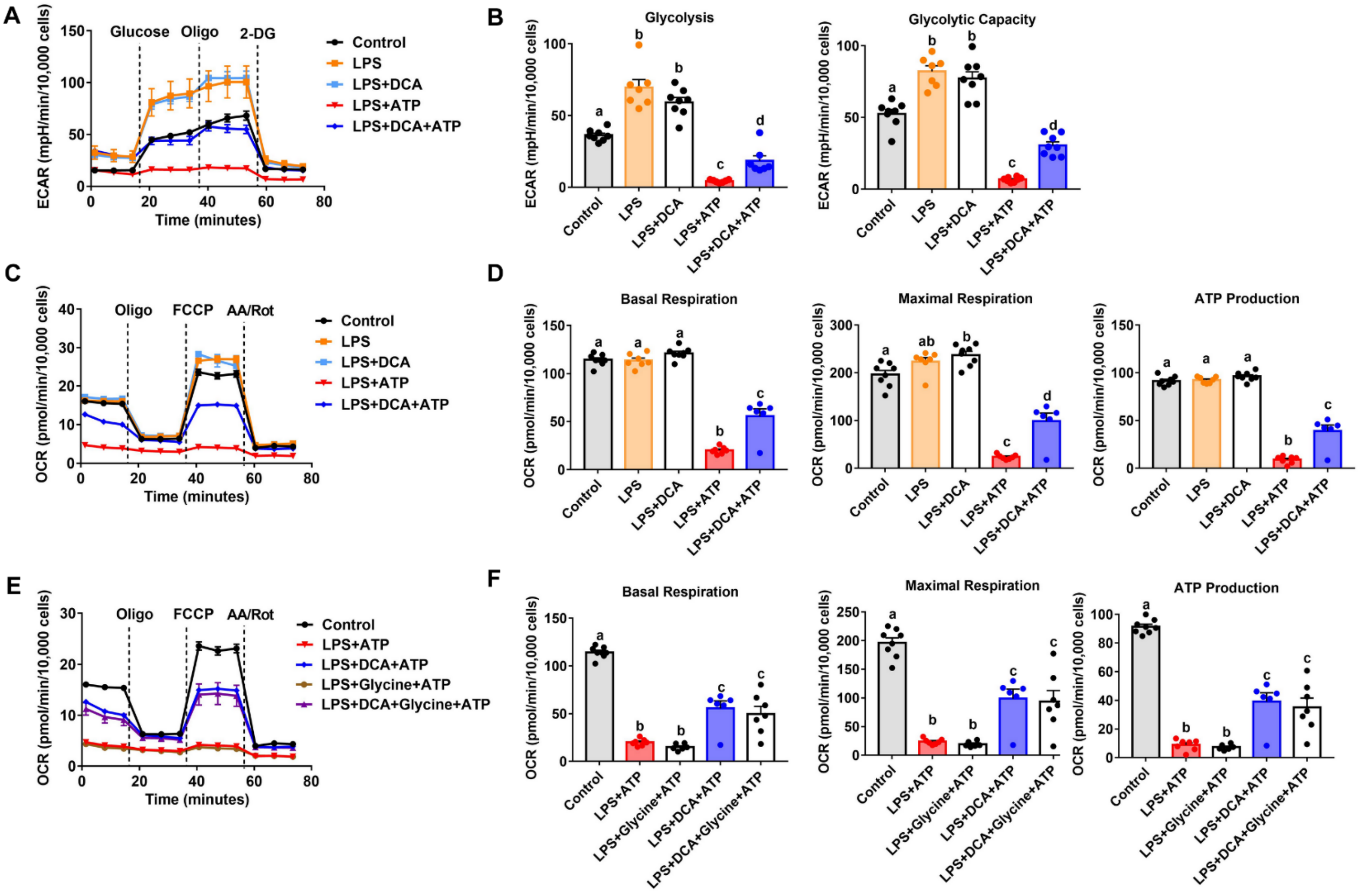


**Figure S2. PDHK inhibition promotes glycolysis and mitochondrial respiration in NLRP3 inflammasome-activated macrophages, Related to Figure 3.**

(**A-B**) Seahorse analysis of extracellular acidification rates (ECAR) in bone marrow-derived macrophages (BMDMs) treated with LPS (300 ng/ml) or LPS (300 ng/ml) + ATP (5 mM) in the presence or absence of DCA (20 mM) for 30 min.

(**C-D**) Seahorse analysis of oxygen consumption rate (OCR) in bone marrow-derived macrophages (BMDMs) treated with LPS (300 ng/ml) or LPS (300 ng/ml) + ATP (5 mM) in the presence or absence of DCA (20 mM) for 30 min.

(**E-F**) Seahorse analysis of oxygen consumption rate (OCR) in bone marrow-derived macrophages (BMDMs) treated with LPS (300 ng/ml) or LPS (300 ng/ml) + ATP (5 mM) in the presence or absence of DCA (20 mM) for 30 min. 5 mM glycine was added 30 min before ATP treatment.

Data are reprehensive of three independent experiments (n=6-8 samples per group). Groups with different letters are significantly different ($p<0.05$); One-way ANOVA with post hoc Tukey's multiple comparisons test.

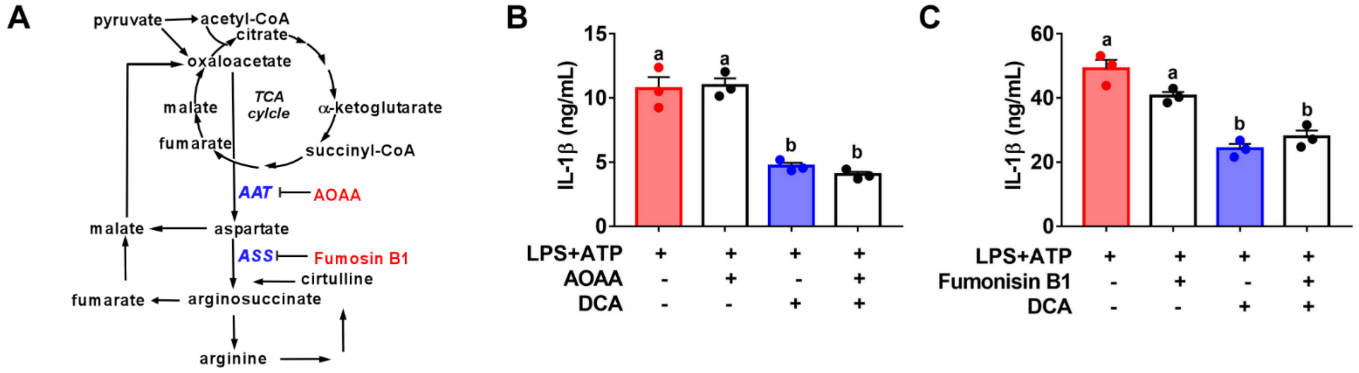


**Figure S3. Inhibition of the malate-aspartate shuttle or aspartate-arginine-succinate shunt does not affect PDHK-regulated NLRP3 inflammasome activation, Related to Figure 4.**

(**A**)The targeting sites of the pharmacological inhibitors are indicated.

(**B-C**) IL-1β secretion from LPS plus ATP-treated BMDMs. BMDMs were stimulated with LPS (300 ng/ml) for 3 h and ATP for 45 min to induce the NLRP3 inflammasome activation. DCA (20 mM) was added after LPS priming. AOAA (1 mM) or fumonisin B1 (15 μM) were added 30 min before ATP treatment.

Data are reprehensive of three independent experiments (n=3-4 samples per group). Groups with different letters are significantly different ($p<0.05$); One-way ANOVA with post hoc Tukey's multiple comparisons test.

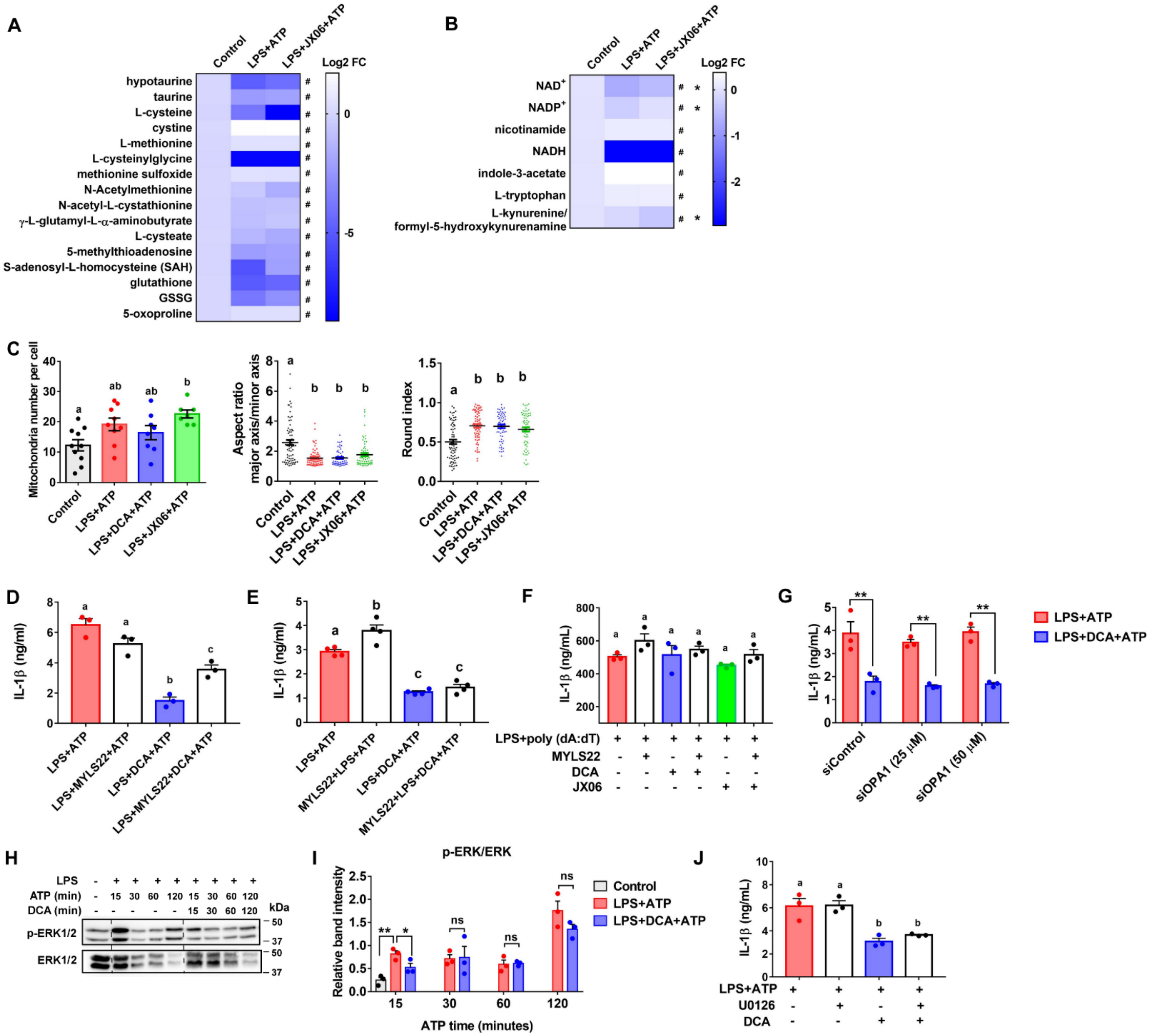


**Figure S4. Redox and mitochondrial remodeling in macrophages treated with or without PDHK inhibitors, Related to Figure 6.**

(**A**) Heatmap of the intermediates in the methionine-cysteine-glutathione pathway analyzed by untargeted metabolomics analysis. LPS-primed bone marrow-derived macrophages (BMDMs) were treated with or without ATP plus JX06 for 45 min.

(**B**) Heatmap of the metabolites in the NAD+ metabolic pathways by untargeted metabolomics analysis. LPS-primed macrophages were treated with or without ATP plus JX06 for 45 min.

(**C**) Mitochondrial number and cristae analysis using transmission electron microscopy. BMDMs were primed with 300 ng/ml LPS for 3 h, followed by 5 mM ATP in the presence or absence of 20 mM DCA or 10 μM JX06 for 45 min.

(**D**) IL-1β level in the culture supernatant from LPS plus ATP-treated BMDMs. OPA1 inhibitor MYLS22 (50 μM) was added 30 min before ATP treatment.

(**E**) IL-1β level in the culture supernatant from LPS plus ATP-treated BMDMs. OPA1 inhibitor MYLS22 (50 μM) was added 30 min before LPS priming.

(**F**) IL-1β level in the culture supernatant from LPS plus poly (dA:dT)-treated BMDMs. BMDMs were primed with 300 ng/ml LPS for 3 h, followed by 2.5 μg/ml poly (dA:dT) in the presence or absence of 20 mM DCA or 10 μM JX06 for 7 h. OPA1 inhibitor MYLS22 (50 μM) was added 30 min before poly (dA:dT) treatment.

OPA1 inhibitor MYLS22 (50 μM) was added 30 min before poly (dA:dT).

(**G)** IL-1β level in the culture supernatant from OPA1 silenced PMs treated with or without LPS plus ATP in the presence or absence of DCA. PMs were treated with 25-50 μM control and OPA1 siRNAs for 72 h before LPS and ATP stimulation.

(**H-I**) LPS-primed BMDMs were stimulated with 5 mM ATP in the presence or absence of 20 mM DCA for 0-120 min. Erk1/2 phosphorylation in macrophages was measured by immunoblotting analysis (E), and the average of the densitometry of the three independent experiments was calculated (F).

(**J**) IL-1β secretion from LPS plus ATP and DCA-treated BMDMs. MEK/Erk1/2 inhibitor U0126 (25 μM) was added 30 min before ATP.

Data are reprehensive of three independent experiments (n=3-4 samples per group). Figures S4A and S4B, $^{\#}$: p<0.05, between LPS+ATP and control groups. *: p<0.05, between LPS+ATP and LPS+JX06+ATP groups. Figures S4C-S4G and S4J, groups with different letters are significantly different (p<0.05). Figure S4I, *: p<0.05, **: p<0.01, ns: non-significant, between groups. One-way ANOVA with post hoc Tukey's multiple comparisons test (Figures S4A-4F and S4J). Unpaired, two-tailed student's T-test (Figures S4G and S4I).
